# Modelling the future distribution of rare bryophytes in Scotland: is inclusion of habitat loss important?

**DOI:** 10.1101/2021.08.30.458156

**Authors:** Anna Ferretto, Pete Smith, David Genney, Robin Matthews, Rob Brooker

## Abstract

Species distribution models (SDMs) have been widely used to predict species ranges and their future distribution under climate change scenarios. In this study we applied Maxent, one of the most used SDMs, to project the distribution of some rare bryophyte species in Scotland in the 2050s. Most of these species are strongly linked to the blanket bog habitat, which is threatened by climate change in the near future. To assess the extent to which changes in habitat distribution leads to a different modelled distribution of the selected bryophytes, blanket bog distribution was included in the model as one of the explanatory variables for some species, and Maxent was run for three 2050s scenarios: once with the current blanket bog distribution and two other runs using the blanket bog distribution derived from two bioclimatic models (Lindsay modified and Blanket Bog Tree model) under the same climate change scenario. For seven out of nine of our studied bryophyte species, the modelled distribution in Scotland was predicted to decline, with some species retreating towards the north-west and other species almost disappearing. When the change in blanket bog distribution was also accounted for, further areas in the north/centre east of Scotland and in the south were predicted to be unfavourable for many of the species considered. Our findings suggest that when modelling species distributions, habitat distribution also needs to be considered, especially when there is a strong relationship between the species and a particular habitat.

## 1 Introduction

As a consequence of climate change, many plant species are being, and are going to be, challenged by unfavourable climatic conditions. In contrast, some may take advantage of increases in the distribution of suitable environmental conditions to increase their overall range, and for others the location of suitable conditions may shift (Berry et al., 2002, Bakkenes et al., 2002, Harrison et al., 2006). In order to plan effective conservation strategies and/or measures to mitigate climate change impacts on species, knowing trends in the distribution of species is necessary information. Species distribution models (SDMs) have been conceived for this, and many other purposes, and are one of the most commonly used tools in this context (Araújo et al., 2011, Hu et al., 2015, Zhang et al., 2019). SDMs define the potential distribution of species by considering their environmental constraints, and one of their applications involves exploration of future scenarios. Although they have some limitations and uncertainties, they have already proved to be a valuable tool for making decisions on conservation issues (Guisan et al., 2012, Gastòn et al., 2014).

The debate around the reliability of SDMs, especially when used to assess future scenarios, has always been lively, with a wide spectrum of suggestions for their improvement, such as the inclusion of non-climatic variables (Stanton et al., 2012; Higa et al., 2013), how to deal with new climatic conditions for which the model has not been trained (Ellis et al., 2014), how to find a balance between excessively complex or simple models (Warren & Siefert, 2011) or how to deal with presence-only data and the consequent selection of pseudo absence data (Iturbide et al., 2018, Phillips et al., 2009, Barbet-Massin et al., 2012).

A discussion that is still not well elaborate, involves how the changing distribution of the main habitat can influence the future distribution of the species. Future scenarios of species distributions are usually obtained accounting for changes in the climatic variables, but the habitat the species rely on – when included - is usually held constant. Pearson et al., (2004) included land cover when modelling the distribution of four plant species in Britain, but for their future projections they kept the land cover constant, changing only the climatic variables. For species that are strongly linked to a specific habitat this might lead to an overestimation of their future distribution, particularly if the habitat is under pressure from climatic and/or non-climatic factors. Stanton et al. (2012) have also pointed out the importance of including non-climatic variables such as vegetation cover and soil type when modelling species distributions, and they argue that, when available, future projections of these variables should be used. Berry et al. (2002) modelled the future distribution of a sample of British and Irish species from different habitats according to their bioclimatic needs and then associated these species to their habitat, trying to infer the future distribution of the habitat from the distribution of some of their key species. Although in this example the issue is considered from the opposite perspective, it underlines the strong link between species and habitat distribution. The same argument has been applied by Oke & Hager (2017), who modelled *Sphagnum* species future distribution in north America to infer *Sphagnum* peatland distribution. The lack of integration of land use change and climate change in studies that investigate species distributions in terrestrial ecosystems has also been pointed out by Sirami et al. (2017), who explained how this could lead to inappropriate management strategies.

In this study, we use Maxent (Maximum entropy model), one of the most widely used SDMs, to model the potential distribution of some rare wetland bryophyte species in Scotland and to predict their future distribution in light of climate change, with and without considering the projected change in the distribution of blanket bog, one of their key habitats. The goals of the study are both to predict how climate change can affect the distribution of these rare species in Scotland and to explore if, and under which circumstances, the SDM prediction is modified by the inclusion of the predicted change in the species’ habitat.

UK bryophytes represent 65% of the European bryophyte diversity and many of them are considered Nationally Rare or Nationally Scarce and are the object of conservation programs and/or designated as features of Sites of Special Scientific Interest (SSSI). Their importance, in terms of biodiversity at an international scale, is due to the oceanic climatic influence that results in many species having their global or European headquarters in the UK and Ireland (Bosanquet et al., 2018). Since many bryophytes do not have active mechanisms to protect themselves from desiccation, these species are very responsive to changes in environmental conditions. Hence, they lend themselves to bioclimatic studies and their future is tightly linked to climate change. Bryophytes have already been the objects of this kind of analysis in the UK. Preston et al. (2011) and Preston et al. (2013) have divided liverworts, hornworts and mosses into clusters according to their recorded distribution, each group resulting in species with similar climatic needs. Smart et al. (2010) have modelled ombrotrophic *Sphagnum* species in the UK and their susceptibility to pollution and climate change, finding that by the 2050s their distribution should be stable or decreased, with the most impacted areas being in the Northwest and some areas in the south of Scotland. Ellis (2015) reviewed the conservation status, the distribution and the implication of climate change for UK bryophytes and lichens, highlighting how – in particular for blanket bog species (and more generally for wetland species) – the potential loss of habitat due to climate change should be considered because it could be the major threat to their survival.

Blanket bog is in fact a strongly climate-dependent peatland habitat that forms in the presence of high rainfall and low evapotranspiration, which allow its typical waterlogged condition to be maintained. In Scotland, it currently covers about 23% of the land area (Bruneau & Johnson, 2014), but climatic projections in the UK forecast warmer and wetter winters and warmer and drier summers (Jerkins et al., 2010), conditions that, overall, might hamper blanket bogs’ preservation. According to some bioclimatic models, in the near future a substantial area of blanket bog is going to be under threat in the UK and in Scotland because of climate change (Clark et al.,2010; Gallego-Sala et al., 2010; Ferretto et al., 2019).

In these circumstances, where the habitat is considered at risk, and where the species are strongly related to this habitat, including information about blanket bog projections provides additional realism to model predictions of the future distribution of the species.

## 2 Methods

We used species distribution models (SDMs) to predict the future distribution of some selected bryophyte species in Scotland. We explored if, for species with a high affinity to blanket bog, SDMs might overestimate their future distribution if they do not account for changes in habitat distribution. One of the most used SDMs is Maxent (Maximum entropy model, Phillips et al., 2006) and we applied it to a list of bryophyte species with different levels of affiliation to blanket bog. First, we projected their distribution to the 2050s climate (medium emission scenario). Then, we repeated the projections taking also into account the potential change in blanket bog distribution due to climate change.

### 2.1 Data preparation

#### 2.1.1 Species selection

A list of bryophyte species with different degrees of obligation with blanket bogs was chosen to explore how the change of their main habitat could change their distribution in Scotland. These species were selected because they are of great significance since they are “Nationally Rare” or “Nationally Scarce” species and are used as one of the parameters to identify Sites of Special Scientific Interest (SSSI) in the UK (Bosanquet et al., 2018). An initial list was made with the help of experts, but it was then reduced to those species for which there were at least 30 records.

Species records were downloaded from the UK NBN Atlas (NBN, 2019 – see Supplementary material 1, Appendix 1 for detailed sources of data). Records were “cleaned” in order to have only accepted presence records (those records that have been verified in the NBN Atlas) in the time frame 1980-2018 and with a spatial uncertainty equal to or lower than 1 km. This time frame was chosen in order to get a balance between a correspondence with the climatic data (1981-2010) and an adequate number of records. The spatial uncertainty in the NBN Atlas spans from 1 m to 10 km: the exclusion of records with an uncertainty higher than 1 km was necessary in order to have the same spatial resolution as the climate data, and to define with sufficient confidence which records were found on blanket bog and which outside it.

The final list was: *Campylopus shawii, Cephalozia loitlesbergeri, Cephalozia macrostachya, Pallavicinia lyellii, Sphagnum lindbergii, Sphagnum majus, Sphagnum pulchrum, Sphagnum riparium* and *Sphagnum strictum* (figure 1). In a recent revision of the phylogeny, the liverworts have in fact been reclassified as *Marchantiophytes* but for simplicity we refer to them all here as *Bryophytes*. Table 1 shows the number of observations for each species and how many of them fall within the blanket bog layer (see section 2.1.3) used in this study or outside it. It also describes the habitats where the species can be found, according to the “Atlas od British and Irish Bryophytes” (Blockeel et al., 2014) and to “The liverworts of Britain and Ireland” (Smith, 1991).

**Table 1.** Number of observations for each species (total, on the blanket bog layer and outside the blanket bog layer), their relative frequency on the blanket bog layer and a description of their habitat.

| Species | Number of records | Number of records on blanket bog | Number of records outside blanket bog | Percentage of records on blanket bog | Habitat description |
| --- | --- | --- | --- | --- | --- |
| *Campylopus shawii* | 63 | 41 | 22 | 65.1% | "Most abundant in flushed mires where it can form very extensive stands, but it also occurs on flushed heathy slopes, by bog pools and on ledges on wet crags. A hyperoceanic species, locally frequent in the Outer Hebrides, but with only a small number of sites in mainland Scotland and south-west Ireland." (Blockeel et al., 2014) |
| *Cephalozia loitlesbergeri* | 60 | 33 | 27 | 55.0% | "Growing over *Sphagnum* and *Leucobryum* in bogs and in moorlands, very rarely on humus in wet rocks clefts." (Smith, 1991) |
| *Cephalozia macrostachya* | 58 | 19 | 39 | 32.8% | "On *Sphagnum* in bogs, rarely on wet peat or wet decaying vegetation." (Smith, 1991) |
| *Pallavicinia lyellii* | 47 | 14 | 33 | 29.8% | "*P. lyellii* grows in two different habitats in Britain. In the north and west, it grows among purple moor-grass (*Molinia*) in wet areas on the fringes of bogs, sometimes at the foot of peat cuttings. In contrast, in the south-east it grows at a few scattered sites in woodlands, either on sandstone, damp, sandy soil or leaf litter." (Blockeel et al., 2014) |
| *Sphagnum lindbergii* | 36 | 4 | 32 | 11.1% | "Found in nutrient-poor flushes and springs high in the mountains, where it can be locally common. Also present at low altitude in and around pools on blanket bogs in Sutherland, and in boggy depressions in Shetland." (Blockeel et al., 2014) |
| *Sphagnum majus* | 31 | 24 | 7 | 77.4% | "Found in nutrient-poor to weakly nutrient-enriched pools and wet hollows in bogs. Its close associates are *S. cuspidatum* and *S. denticulatum*, and less frequently *S. fallax*. *S. majus* does not occur submerged, except during floods." (Blockeel et al., 2014) |
| *Sphagnum pulchrum* | 126 | 44 | 82 | 34.9% | "Occurs in pools and hollows in bogs and valley mires, and in acidic flushes. Widespread, but very local; commonest on Dorset heaths." (Blockeel et al., 2014) |
| *Sphagnum riparium* | 41 | 13 | 28 | 31.7% | "Typically found in weakly to moderately base-rich springs, flushes and in poor fens." (Blockeel et al., 2014) |
| *Sphagnum strictum* | 83 | 15 | 68 | 18.1% | "Occurs, often amongst purple moor-grass (*Molinia*) in wet heath, boggy grassland, shallow blanket bog and on very humid, rocky slopes. Almost exclusively western, growing in districts with high rainfall." (Blockeel et al., 2014) |

**Figure 1.**
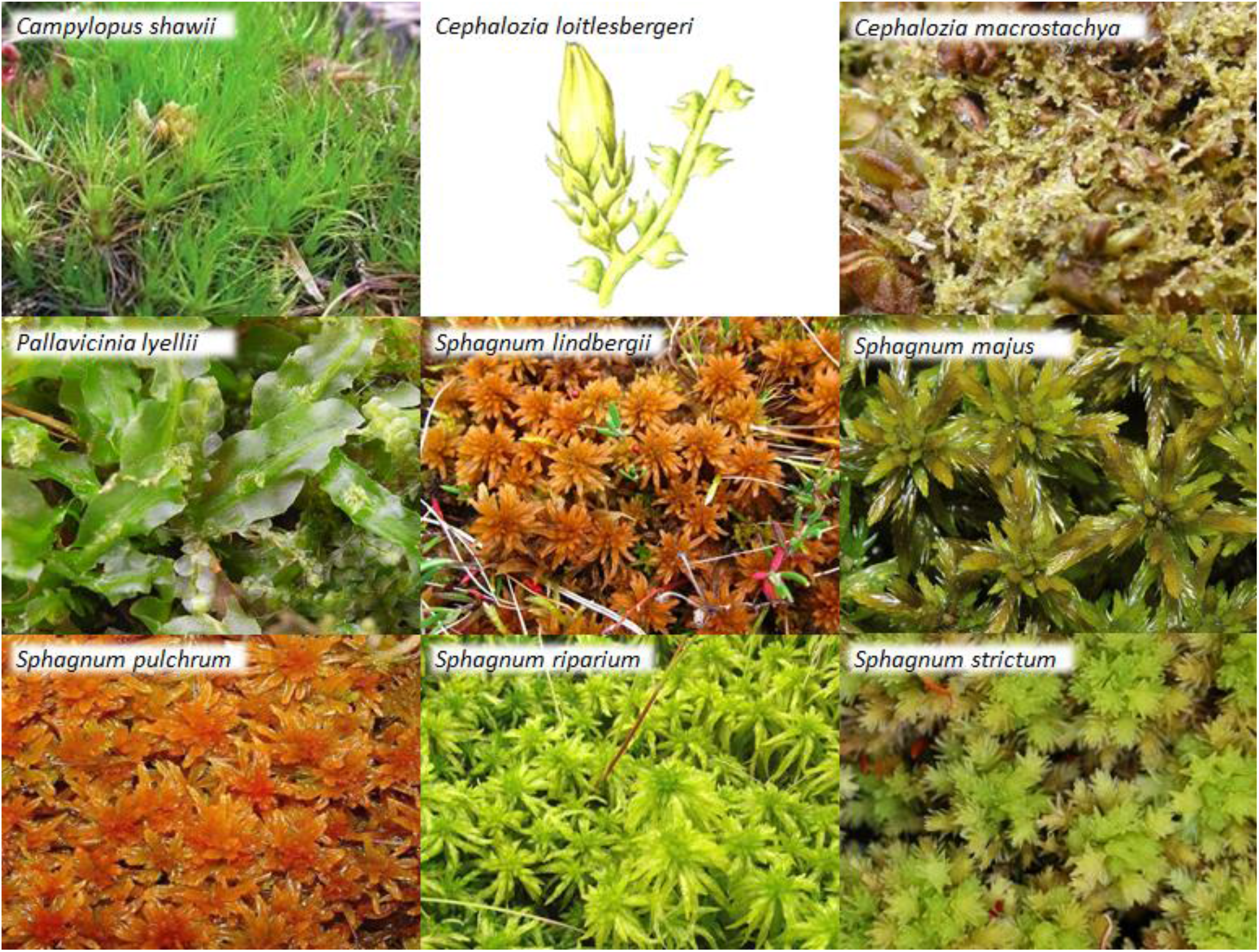
Bryophyte species considered in this study.

#### 2.1.2 Climate data

Climate data were obtained from the HadUK-Grid model that covers the UK at a resolution of 1 km (Hollis et al., 2019). Data are averaged for the period 1981-2010. Annual, monthly and seasonal data are available for mean, minimum and maximum temperature and for total precipitation. To explain variation in species distribution, we used annual mean temperature (°C) and annual total precipitation (mm) and from monthly data we derived values for summer (June to August) precipitation (mm), winter (December to February) precipitation (mm), maximum temperature of the warmest month (°C), minimum temperature of the coldest month (°C) and potential evapotranspiration (PET) (mm).

#### 2.1.3 Blanket bog distribution

A map of current blanket bog distribution in the UK was obtained by extracting the class “bog” from the UK Land Cover Map 2007 at a resolution of 25 m (Morton et al., 2011).

#### 2.1.4 Slope

A Digital Elevation Model (DEM) of the UK, was cut from the NASA’s Shuttle Radar Topography Mission (SRTM) DEM at 90m resolution (Jarvis et al., 2008). From this DEM we calculated the slope. All the data were then processed to get the same coordinate system, extension and resolution (1 km grid). A preliminary analysis was also run including pollution data (NOx, NHx and SOx) but, given the different spatial resolution of these data (5km) compared to the climate data, they introduced further uncertainty and the results (similar for most of the species, especially considering the current distribution) were not reported in the main text (they are included in Supplementary material 1, Appendix 6).

### 2.2 Models choice and settings

Maxent is one of the most commonly used models for species distribution. In particular, it performs better than other models when the sample size is small (Williams et al., 2009) and it can be used with ‘presence only’ data, as was the case for the NBN Atlas species records. The analysis was carried out with the software MAXENT Version 3.4.0 (Phillips et al., 2006). The parameters used to run MaxEnt have been carefully chosen according to what is suggested in the literature, and they are described in the next paragraph and summarised in the ODMAP protocol (Supplementary material 2), following Zurell et al. (2020).

#### 2.2.1 MaxEnt

Using species presence-only locations and a set of environmental predictors at those locations, Maxent finds the distribution that is both most spread out (with the maximum entropy) and within the limits of the environmental predictors. It compares the environment found at locations where the species is present, with the environment found in a sample of background locations taken from the study area and where presence is unknown.

For each of the grid cells into which the area is divided, Maxent returns the probability - given the species is present - that it is found in that grid. This corresponds to the ‘raw output’ of Maxent. The program can also return the ‘logistic output’, which is the estimate of the probability that a species is present, given the environment. This is more interpretable and always has values between 0 and 1. The main problem with the logistic output is that it makes an assumption concerning the probability of presence (π) at sites with typical condition for the species. The default value for π is 0.5, which means that the species is present in half of all the possible locations. While Elith et al. (2011) suggest that the logistic output with the default value gives a better result than the raw output, Merow et al. (2013) do not recommend using the logistic output without justifying the choice of π. For a better interpretability of the results, we decided to use the logistic output, but since we are modelling rare species, we arbitrarily chose a lower value for π (0.1).

The model applies some transformations of the predictors (called features); users can choose between different features (linear, quadratic, product, threshold and hinge features for continuous variables, and discrete features for categorical variables) according to the number of records available. As suggested by Phillips & Dudìk (2008), for presence-only data and with more than 15 records, we used the hinge feature.

To avoid overfitting, Maxent uses a regularization parameter (β). The default value for β is 0.5 and it has been tested for a wide range of taxonomic groups (Phillips & Dudìk, 2008). Merow et al. (2013) suggest exploring a range of values for the regularization coefficient. We used the R package “*MaxentVariableSelection*” (Jueterbock et al., 2016) both to find the best value of β and to operate a variable selection. The package fits various combinations of variables and β values and offers two options to select them: the model that maximizes the area under the receiving operating characteristic curve (AUC) and the model that minimizes the Akaike Information Criterium (AIC). AUC measures how the model can discern between classes, while AIC score is a measure of how well the model fits the data, without overfitting it. When Maxent is used to obtain future projections, Warren & Seifert (2011) suggest using the AIC method because it is more transferable to future climate scenarios. We selected the model with the lowest AIC.

Finally, we converted the probability distribution into binary (presence/absence) maps. When possible, it is preferable to use probability maps, because the conversion adds more uncertainty - related to the choice of the threshold - and reduces the information provided (Guillera-Arroita et al., 2015). In this case, we needed to compare the outcomes of three different distributions to assess how the species ranges change as a consequence of climate change (see par. 2.3) and the binary maps were necessary; probability maps can be found in Supplementary material 1, Appendix 2. For each species, we chose the threshold that maximized the sum of sensitivity and specificity (as suggested by Liu et al., 2005) in the training dataset when calculating the present distribution and we used the same thresholds also for the binary conversion of the distribution of the future projections.

Maxent assumes that any location is equally likely to be sampled. Since the sampling effort of the selected bryophyte records was unknown, we used “target group sampling” (TGS) proposed by Phillips et al. (2009) to account for sample bias. This method uses presence locations of other species (taxonomically related to the focal species) to estimate sampling, by assuming that in the same survey, the focal species would have been found if the focal species were present in those locations. This allows modification of the background (“biased background”) in order to account for the survey effort. Bystriakova et al. (2012) demonstrated how this approach can increase the predictive ability of Maxent; Syfert et al. (2013) claim that correcting for geographic bias should be done by default when running Maxent. To build our biased background, we used records from all the bryophyte species observed in the UK NBN Atlas (with the same selection made for the focal species about year, spatial uncertainty and acceptance status). For each species, the number of background points was set ten times higher than the number of presences, in accordance with Barbet-Massin et. al (2012).

To evaluate the models, we used the area under the receiving operator curve (AUC), tested with a 5-fold cross validation. Since presence data are opposed to background data (and not to absence data), the AUC in this case is a measure of the ability of the model to discern between presence and background. AUC values span from 0 to 1, with a value of 1 meaning that the model can perfectly classify the predictions and a value of 0.5 meaning that the classification returned by the model is no better than random. We selected models with AUC values > 0.7, as this is a sign of a reasonable discrimination ability (Pearce & Ferrier, 2000). Alongside with the AUC, we used the Boyce index, which is particularly suitable for presence-only data. The Boyce Index can vary between -1 and 1, where values close to zero suggest that the model is not different from a chance model, negative values suggest a wrong model, where areas with presences are more frequent predicted as not suitable, and positive values suggest a strong model, where the predictions coincide with the presence’s distribution in the evaluation dataset (Boyce et al., 2002; Hirzel et al., 2006). In order to help understand the relative importance of the single variables used in the models, Maxent returns the estimates of their relative contributions. We also ran the jackknife test, which excludes each variable in turn and creates a model with the remaining variables, and a model with each variable used by itself. Comparing them with the model with the full set of variables allows us to understand which variables contribute the most to increase the fit and which variables carry more information by themselves.

### 2.3 Predictions

One of the problems in using SDMs for future scenarios is that the models might have to deal with climate variables that are outside the ranges seen in the training data. In order to avoid this, the model was fitted and tested for the UK and projected only for Scotland. Scotland is in fact expected to warm in the near future, and to experience a climate that will be comparable with the current climate of some areas of southern England. In this way, we avoid challenging the model with untested conditions. The same approach was taken by Ellis et al. (2016) for modelling lichen species distribution in the north of the UK.

For the climatic variables, the climate projections at 25 km2 resolution were downloaded from the UKCP09 website for the medium emission scenario in the 2050s (Hadley Centre for Climate Prediction and Research, 2017). We downscaled the data to 1 km resolution and created a mask on Scotland. More recent and finer projections were available (UKCP18), but only for the high emission scenario. Besides this, we wanted to maintain consistency with data used in Ferretto et al. (2019), which was used to estimate future blanket bog distribution.

Slope values were not changed for forward projections.

To check if blanket bog distribution may have an impact on the projected distribution of the species, we ran the model twice: the first time keeping the current blanket bog distribution (with a mask on Scotland only), and the second time with the projected blanket bog distribution obtained from Ferretto et al. (2019), calculated for the 2050s under the medium emission scenario, with two models (Blanket Bog Tree model -BT- and Lindsay Modified model - LM). These distributions are shown in figure 2, while the equations of the two models are reported in Appendix 3.

**Figure 2.**
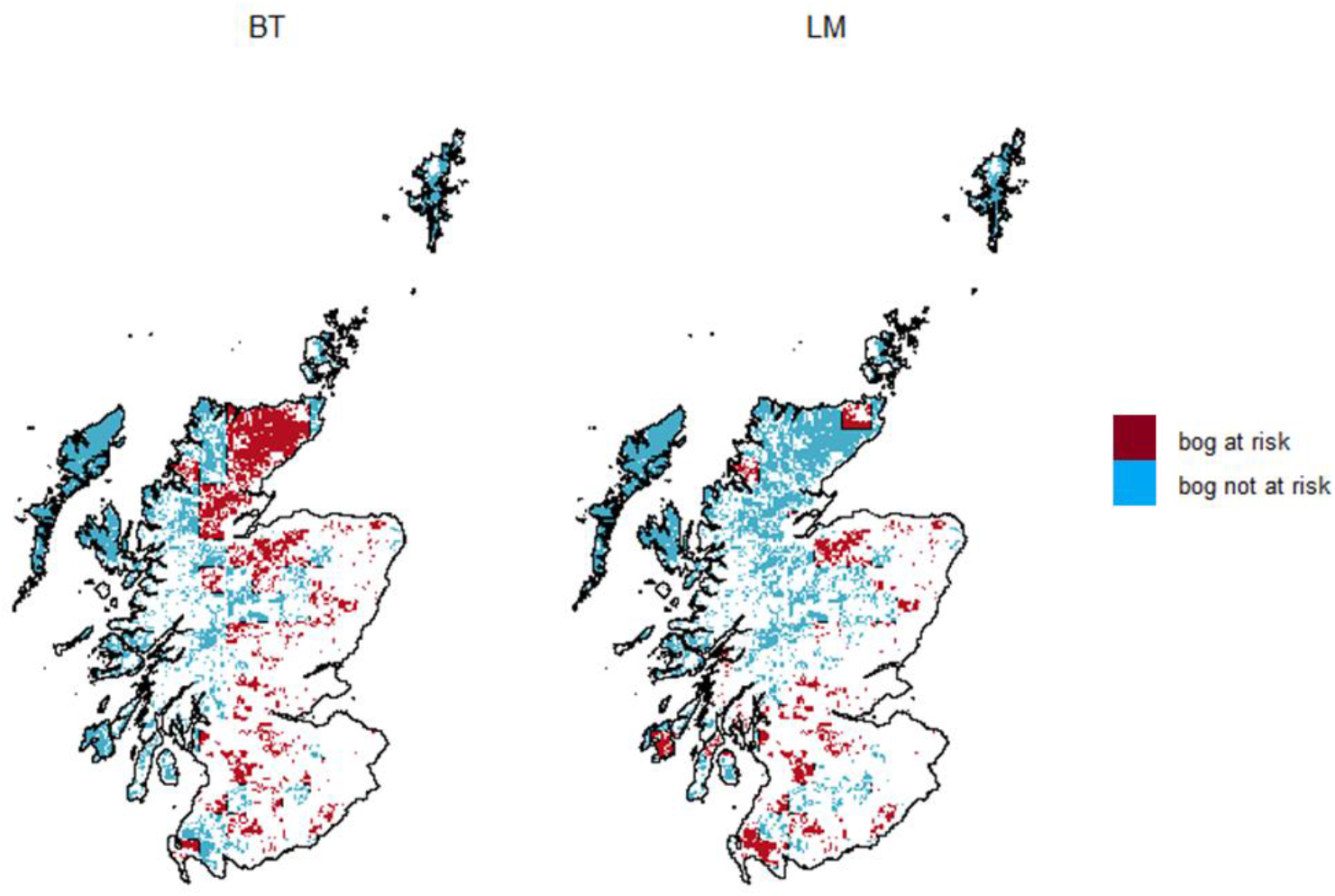
Blanket bog future (2050s) predicted distribution, medium emission scenario obtained using two different bioclimatic envelope models: Blanket bog Tree model (BT) on the left and Lindsay Modified model (LM) on the right (adapted from Ferretto et al., 2019). Blanket bog not threatened by climate change in blue. In red, blanket bog at risk of loss due to climate change.

### 2.4 Comparison

The predictions obtained for each species in the previous step were compared to evaluate how the loss of habitat can affect species distribution. The difference between the two projections (with a fixed blanket bog layer and with a projection of the blanket bog layer that takes into account climate change) were computed.

## 3 Results

### 3.1 Model performance

For all the species, Maxent performed well, with test AUC values averaged for the five replicates always > 0.7 and > 0.8 for all the species except that for *Cephalozia macrostachya* and *Pallavicinia lyellii* and the Boyce Index always positive and with the lowest value being 0.799 (table 2).

**Table 2.** Test AUC values and their standard deviation and Boyce Index for each species.

| Species | Average test AUC | Standard deviation | Boyce Index |
| --- | --- | --- | --- |
| *Campylopus shawii* | 0.943 | 0.008 | 0.987 |
| *Cephalozia loitlesbergeri* | 0.870 | 0.007 | 0.827 |
| *Cephalozia macrostachya* | 0.775 | 0.077 | 0.956 |
| *Pallavicinia lyellii* | 0.743 | 0.071 | 0.940 |
| *Sphagnum lindbergii* | 0.953 | 0.014 | 0.983 |
| *Sphagnum majus* | 0.943 | 0.017 | 0.877 |
| *Sphagnum pulchrum* | 0.926 | 0.022 | 0.799 |
| *Sphagnum riparium* | 0.897 | 0.040 | 0.928 |
| *Sphagnum strictum* | 0.917 | 0.016 | 0.931 |

The variable selection process led to the choice of the variables and of the β values reported in table 3. These were the combinations of β values and variables that minimized the AIC score.

**Table 3.**
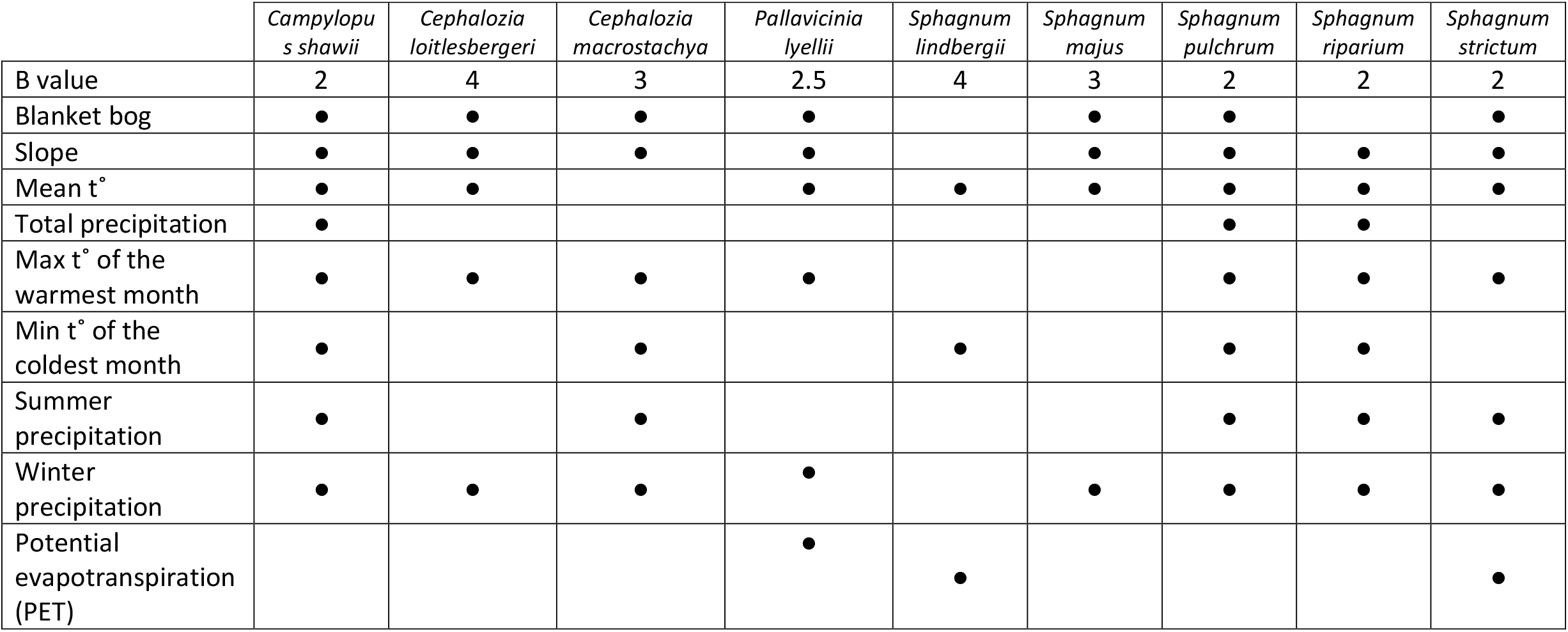
Regularization parameters (β values) and variables used to run Maxent for each species.

|  | *Campylopus shawii* | *Cephalozia loitlesbergeri* | *Cephalozia macrostachya* | *Pallavicinia lyellii* | *Sphagnum lindbergii* | *Sphagnum majus* | *Sphagnum pulchrum* | *Sphagnum riparium* | *Sphagnum strictum* |
| --- | --- | --- | --- | --- | --- | --- | --- | --- | --- |
| B value | 2 | 4 | 3 | 2.5 | 4 | 3 | 2 | 2 | 2 |
| Blanket bog | • | • | • | • |  | • | • |  | • |
| Slope | • | • | • | • |  | • | • | • | • |
| Mean t° | • | • |  | • | • | • | • | • | • |
| Total precipitation | • |  |  |  |  |  | • | • |  |
| Max t° of the warmest month | • | • | • | • |  |  | • | • | • |
| Min t° of the coldest month | • |  | • |  | • |  | • | • |  |
| Summer precipitation | • |  | • |  |  |  | • | • | • |
| Winter precipitation | • | • | • | • |  | • | • | • | • |
| Potential evapotranspiration (PET) |  |  |  | • | • |  |  |  | • |

### 3.2 Current distribution

The current distribution predicted by Maxent is shown in figure 3. The presence/absence maps were obtained from the probability distribution maps (Appendix 2 in Supplementary material 1) by applying the threshold that maximized the sum of sensitivity and specificity in the training dataset (table 4).

**Table 4.**
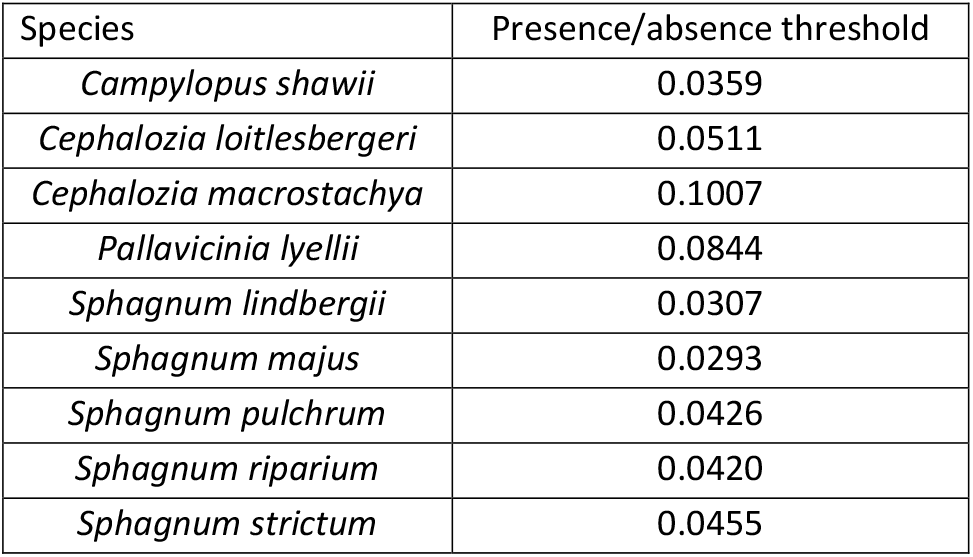
Maxent thresholds for the presence/absence conversion, obtained maximizing the sum of sensitivity and specificity in the training dataset.

| Species | Presence/absence threshold |
| --- | --- |
| *Campylopus shawii* | 0.0359 |
| *Cephalozia loitlesbergeri* | 0.0511 |
| *Cephalozia macrostachya* | 0.1007 |
| *Pallavicinia lyellii* | 0.0844 |
| *Sphagnum lindbergii* | 0.0307 |
| *Sphagnum majus* | 0.0293 |
| *Sphagnum pulchrum* | 0.0426 |
| *Sphagnum riparium* | 0.0420 |
| *Sphagnum strictum* | 0.0455 |

**Figure 3.**
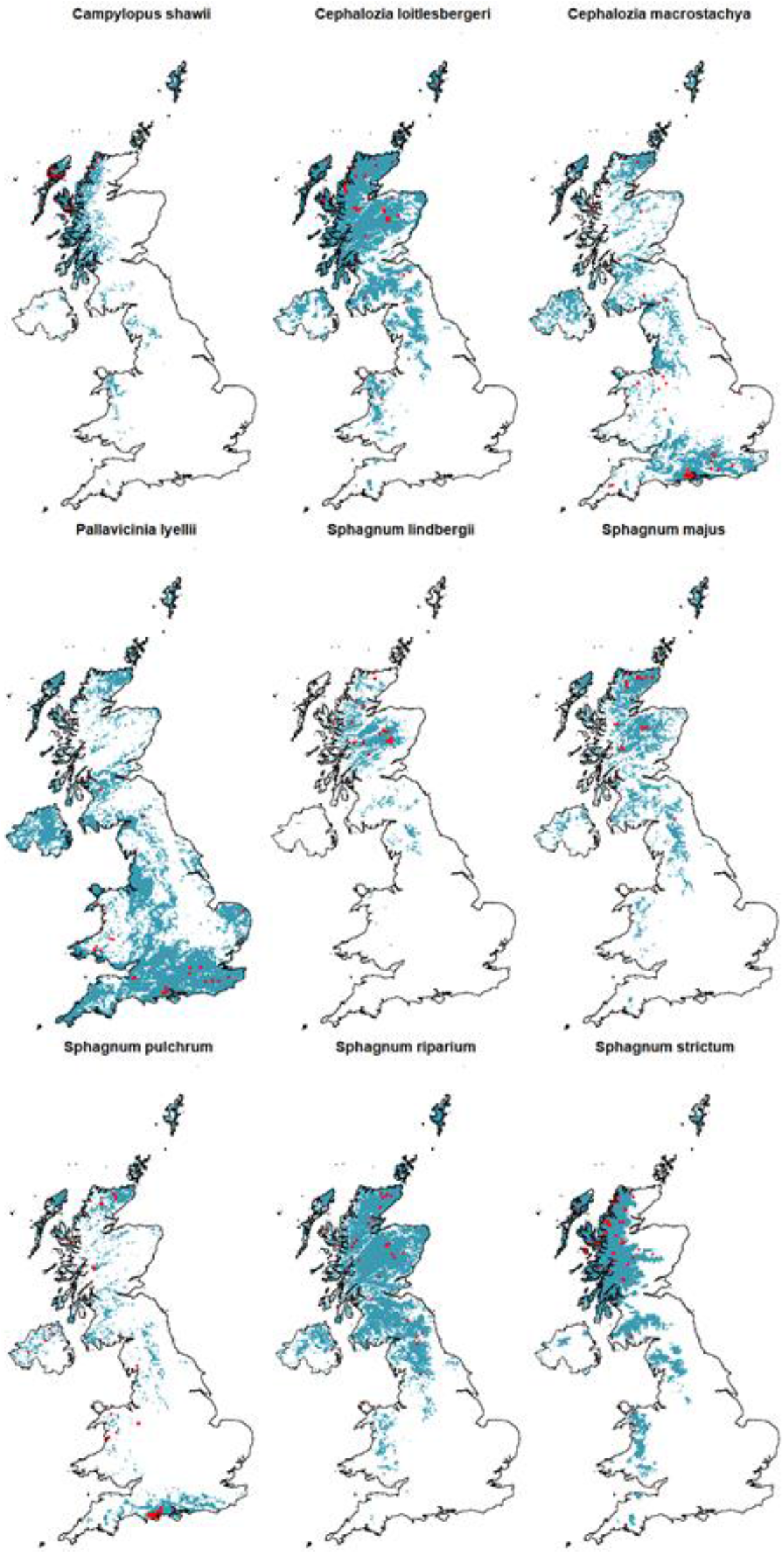
Current species distribution modelled with Maxent in the UK (blue) and observed records (red points).

For most of the species, the distribution predicted by Maxent is primarily located in Scotland, Wales and Northern Ireland, with only three species (*Cephalozia macrostachya, Pallavicinia lyellii* and *Sphagnum pulchrum*) that are widespread in the warmer southern England. In particular, the modelled distributions of *Campylopus shawii* and *Sphagnum strictum* cover only the humid west of Great Britain, mostly in Scotland, including all the islands, and to lesser extent in Wales, northern England and very few patches in Cornwall. A small area of the northern coast of Northern Ireland completes the picture. *Cephalozia loitlesbergeri, Sphagnum riparium, Sphagnum lindbergii* and *Sphagnum majus* form another group of species with a similar modelled distribution, occupying the northern part of Scotland (mostly the west and the Highlands but also some areas of the east coast) and parts of southern Scotland. They are also present in the central part of northern England, in Wales and in the north coast of Northern Ireland. Despite the similar pattern, the presence in these areas is more marked for *Cephalozia loitlesbergeri* and *Sphagnum riparium*, which have a widespread distribution almost in the whole Scotland, compared to *Sphagnum majus* and *Sphagnum lindbergii*, the last one being absent from the Scottish islands, the east coast of Scotland and with only few patches in Wales and Northern Ireland. *Cephalozia macrostachya, Pallavicinia lyellii* and *Sphagnum pulchrum* have the widest north-south and east-west modelled range, with presences from the westernmost corner of Cornwall to the easternmost coast of Kent, and from the southern coast of England to the north coast of Scotland and Shetland Islands. Interestingly, they are also the only species with a notable presence in the central belt in Scotland. Despite their widespread modelled distribution, they are not prevalent in the north-west of Scotland and in the central area of England.

### 3.3 Future distribution

#### 3.3.1 With the same blanket bog distribution in the future

We projected species distributions into the future (2050s), with climate variables changed to account for the climate change (according to the medium emission scenario), while slope and blanket bog distribution were kept at their current values.

When projected into the future, seven species see a decrease in their predicted suitable area in Scotland and two species are predicted to increase their distribution (second column of table 5 and figure 4).

**Table 5.** Change in the predicted future distribution (2050s) compared to the current modelled distribution for each species, obtained considering only climate variables or climate variables and predicted blanket bog distribution according to BT and LM models.

| Species | % of change | % of change BT<br>bog distribution | % of change LM<br>bog distribution |
| --- | --- | --- | --- |
| *Campylopus shawii* | -34.1 | -37.2 | -35.3 |
| *Cephalozia loitlesbergeri* | -53.9 | -60.4 | -55.6 |
| *Cephalozia macrostachya* | +9.5 | -14.6 | -3.4 |
| *Pallavicinia lyellii* | +83.1 | +63.5 | +75.0 |
| *Sphagnum lindbergii* | -100.0 | / | / |
| *Sphagnum majus* | -98.0 | -98.8 | -98.4 |
| *Sphagnum pulchrum* | -4.4 | -36.6 | -23.9 |
| *Sphagnum riparium* | -90.5 | / | / |
| *Sphagnum strictum* | -33.8 | -33.3 | -33.8 |

**Figure 4.**
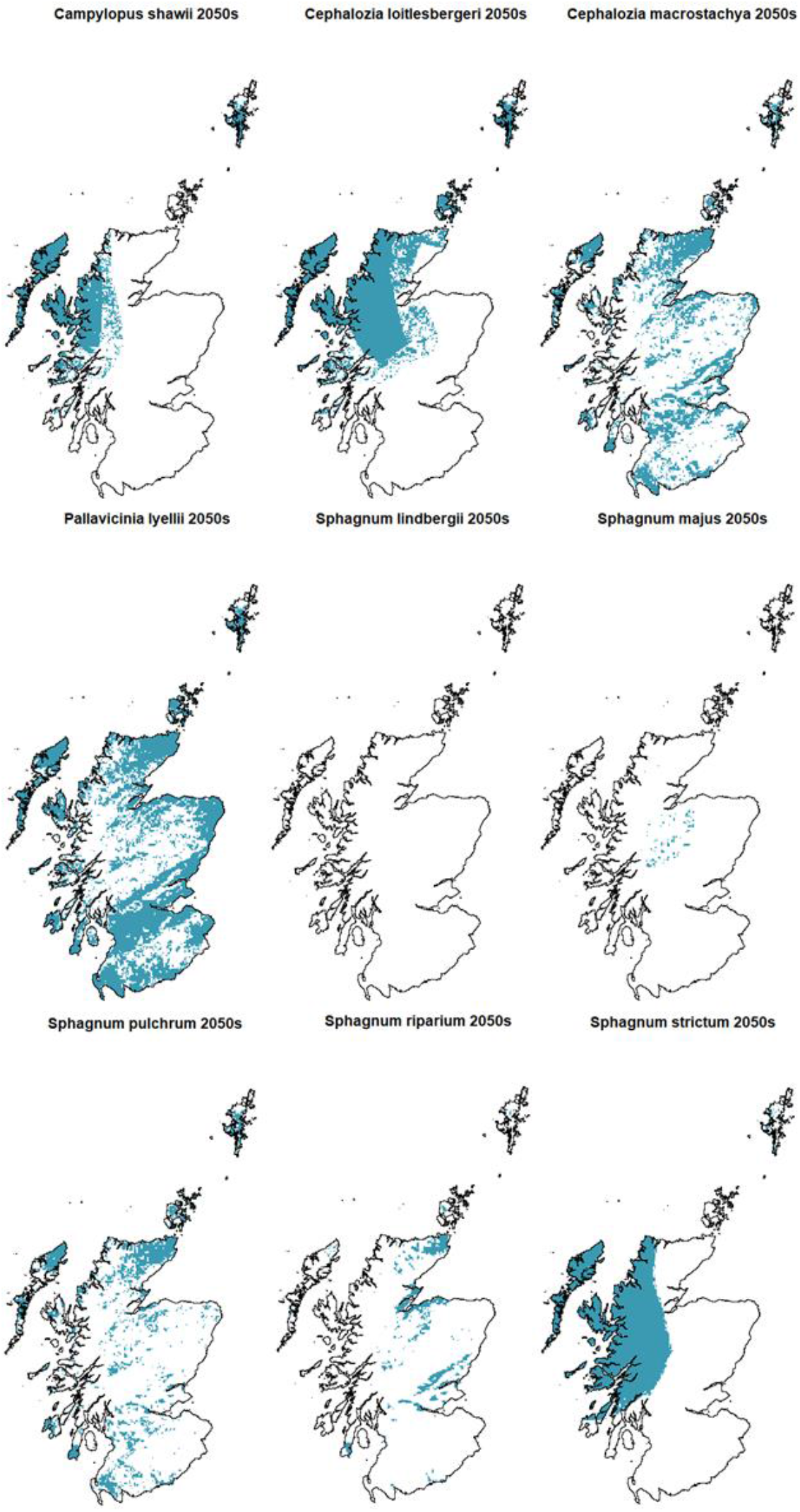
2050s species distribution predicted with Maxent in Scotland (medium emission scenario) obtained considering only changes in the climatic variables.

The species predicted to increase their distribution are *Cephalozia macrostachya* (+9.58%) and *Pallavicinia lyellii* (+83.1%). This increase is happening around areas already favourable to the species. All the other species are predicted to diminish their favourable areas, from *Sphagnum pulchrum* (−4.4%) to *Sphagnum lindbergii*, which is predicted to disappear from Scotland. According to Maxent predictions, the distribution of *Sphagnum pulchrum* is predicted to remain very similar to the current predicted distribution. *Campylopus shawii and Sphagnum strictum*, which have a similar distribution, are both predicted to retreat towards the northwest, losing around 30% of their current favourable area, with a neat longitudinal threshold (in the middle of the Highlands for *Sphagnum strictum* and in the western highlands for *Campylopus shawii*) and only the Shetland Islands further east than this line. *Cephalozia loitlesbergeri* is predicted to retreat towards the northwest and the islands (−53.9%), losing all the favourable areas in the South and a substantial area in the East. Finally, *Sphagnum majus* and *Sphagnum riparium* are predicted to lose almost their entire area (−98.0% and -90.5%), with Sphagnum majus that keeps only a small portion in the middle of the Highlands and *Sphagnum riparium* which maintains its presence in the islands, in the northeast and in some areas of the east central Scotland.

#### 3.3.2 Changing the blanket bog layer

We also tested whether accounting for a change in blanket bog distribution might affect future species distributions. The variable selection process ran for each species excluded the blanket bog layer for *Sphagnum lindbergii* and *Sphagnum riparium* (Table 3), which were hence excluded from this further step. Both these species are in fact not strictly blanket bog species: *Sphagnum lindbergii* is found in blanket bogs near pools in Sutherland, but its main habitat are oligotrophic flushes and springs, while *Sphagnum riparium* is a poor fen species (Blockeel et al., 2014). Only 11.1% of the observed records of *Sphagnum lindbergii* are within the blanket bog layer, while for *Sphagnum riparium* the percentage is higher (31.7%).

When the change in blanket bog distribution is taken into account, in terms of total area, the major differences are predicted for *Pallavicinia lyellii, Cephalozia macrostachya* and *Sphagnum pulchrum*. For *Pallavicinia lyellii*, there is still a gain of suitable area (+63.5% and +75% with the BT and LM distribution respectively), to lesser extent than in the initial model (+83%). *Cephalozia macrostachya*, that according to the initial model will increase its suitable area of 9.4%, with the BT and LM blanket bog distribution is instead predicted to lose 14.6% and 3.4% of its distribution. For *Sphagnum pulchrum*, whose distribution was predicted to decrease of 4.4%, the loss of suitable area is much higher when the BT and LM blanket bog distributions are used (−36.6% and -23.9%). In particular, for these three species, the greatest losses compared to the initial model are in the Central Belt, in the south and in the east of Scotland (for both the BT and LM distribution) and especially marked in the northeast when the BT distribution is used. For all the other species, the differences between the initial model and the models ran with the BT and LM blanket bog distributions are much less marked. *Cephalozia loitlesbergeri* and *Sphagnum majus* lose some favourable areas in the eastern border of their distribution. This loss is more evident for *Cephalozia loitlesbergeri* than for *Campylopus shawii* and with the BT distribution than with the LM distribution. *Sphagnum majus*, which was already predicted to lose most of its extension (−98%), with the BT and LM distribution is projected to lose a further area when the BT and LM blanket bog distributions are used. No differences are detected for *Sphagnum strictum*.

Percentages of gain and losses obtained running the models with the BT and the LM blanket bog distributions are listed in table 5, while figure 5 shows the distributions of the species predicted in this step and the differences with the initial model.

**Figure 5.**
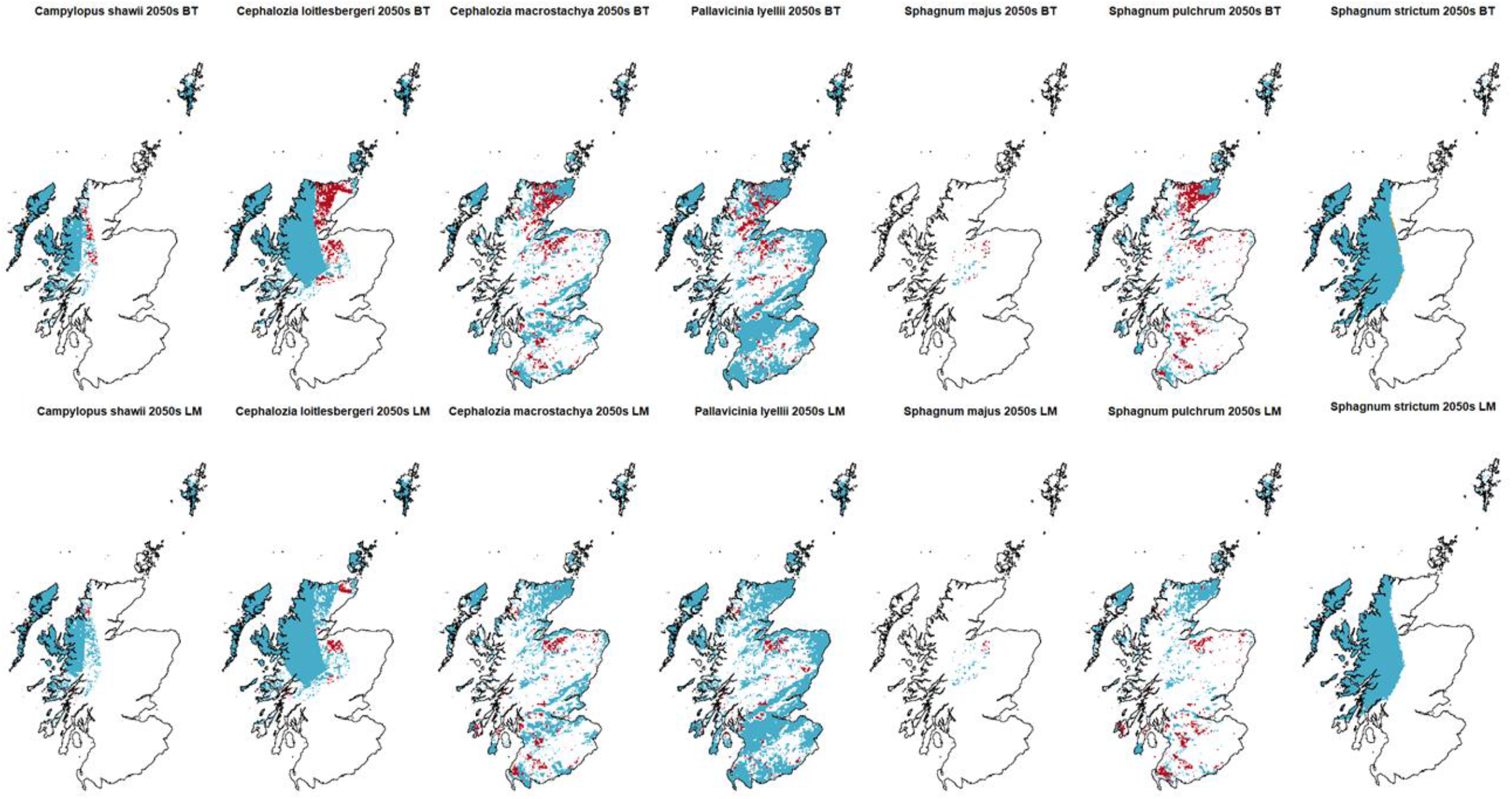
Species distribution modelled by Maxent in the 2050s and using the BT (above) and LM (below) blanket bog distribution. In blue: future distribution as per the initial model (figure 4) that is unaffected by changes in the blanket bog layer. In red: areas that are predicted to be suitable by the initial model but are not suitable if blanket bog modelled change is considered. The very thin yellow strip in Sphagnum strictum, BT distribution, indicates areas not predicted to be suitable by the initial model, but suitable if blanket bog change is considered

**Figure 5.**
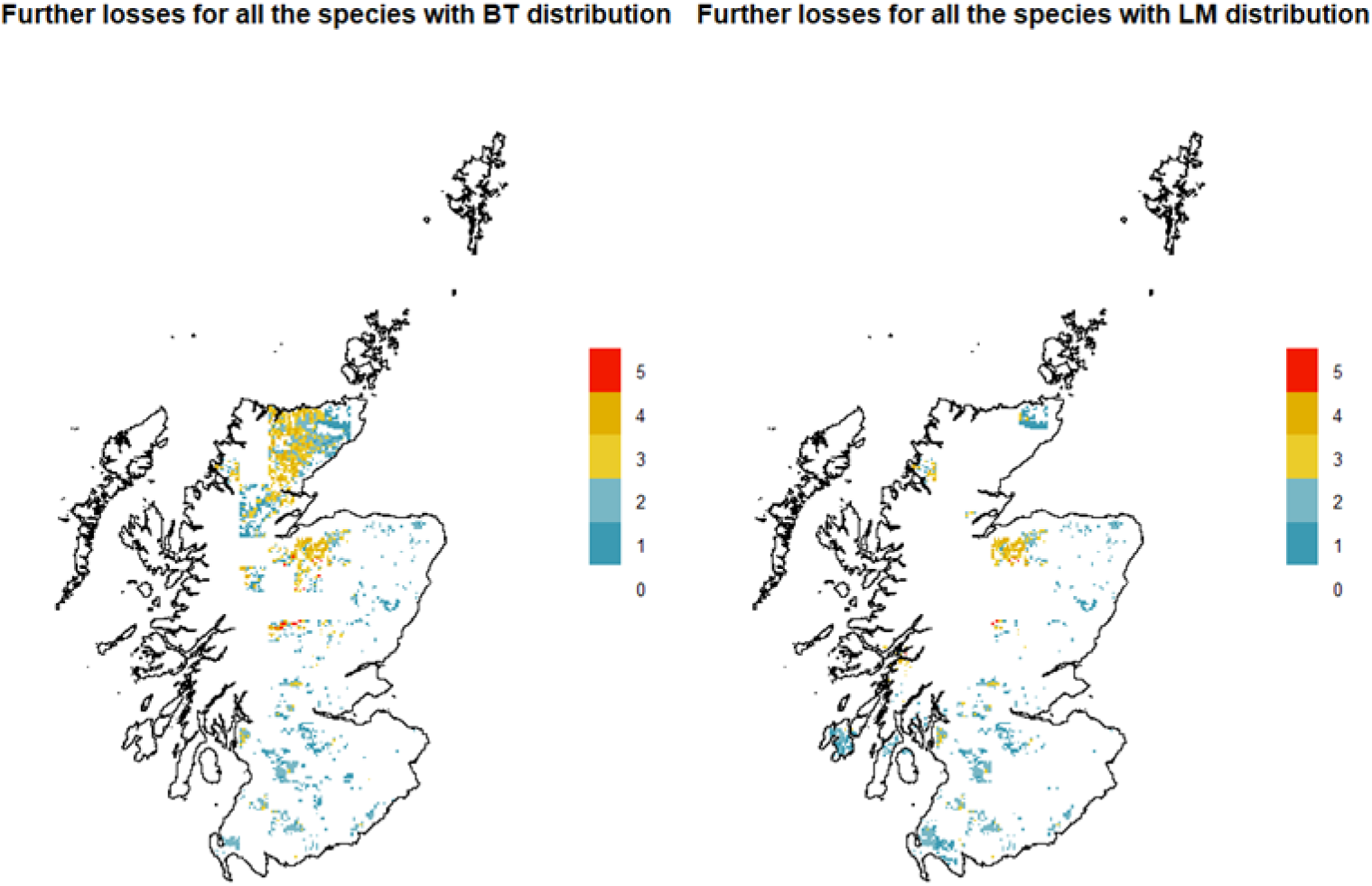
Areas most affected by the change in blanket bog modelled distribution, considering all the species (BT distribution on the left, LM distribution on the right). The legend shows the number of species involved.

Considering all species, areas where there are more losses between the initial model and the model with BT distribution are located in the northeast of Scotland, at the border between the Highlands and Moray, in Perthshire and some areas in the south. The same general pattern is seen with LM bog distribution, but without reductions in the northeast (figure 6).

### 3.4 Importance of the variables

For the three species that are predicted to lose almost entirely their suitable area (*Sphagnum lindbergii, Sphagnum majus* and *Sphagnum riparium*), the jacknife test reveals that mean temperature is the variable with the highest gain when used alone (which means that it is the variable that carries most of the information) and for *Sphagnum lindbergii* and *Sphagnum majus* it leads to the highest loss of fit in the model if it is removed (which means that has the most information that is not present in the other variables). It is also the variable with the highest contribution to the model for *Sphagnum lindbergi* (98.4%) and for *Sphagnum riparium* (78.1%). The species with the greatest difference in the predicted distribution between the initial model and the models run with the BT and LM blanket bog distributions are *Pallavicinia lyellii, Sphagnum pulchrum* and *Cephalozia macrostachya*. In these three species, the variable blanket bog has a great importance, being either the variable that gives the highest fit if used in isolation (for *Cephalozia machrostachya*) or the variable that, if omitted, generates the highest loss of fit in the model (for *Pallavicinia lyellii* and *Sphagnum pulchrum*). For *Pallavicinia lyellii* and *Cephalozia machrostachya*, it is also the variable that, after the slope, contributes the most to the model (22.9% and 31.7% respectively). Blanket bog is also the most important variable for *Sphagnum majus* (45.1%), although in this case there is not a great difference between the initial model and the model run with the BT and LM distribution. In this case, however, the initial model already predicted a 98% loss of suitable area, and it is hence difficult to evaluate the contribution of the blanket bog variable for the remaining 2%. In *Campylopus shawii* and *Cephalozia loitlesbergeri*, the maximum temperature of the warmest month is both the variable that alone carries the most information, and the variable that, if removed, leads to the highest loss of fit of the model. It is also the variable with the greatest contribution to the model for these species. Finally, winter precipitation is the most important variable for *Sphagnum strictum* under all the points. Although blanket bog is inserted in the model of *Sphagnum strictum*, its contribution to the model is null (0.2%), and this explains the fact that there is no difference between the initial model and the models with BT and LM blanket bog distribution. The contribution of each variable and the results of the jackknife test for each species are reported in Appendices 4 and 5 (Supplementary material 1).

## 4. Discussion

The goals of the study were both to model the future distribution of some rare bryophyte species in Scotland and to compare the distributions obtained with and without considering predicted changes in their main habitat due to climate change.

### 4.1 Model performance and uncertainties

The model performed well for all the species, with excellent values of AUC (that, in the case of presence-only data, indicate the ability to discern between presence and background data) and always positive values of the Boyce Index. The only models with an AUC value lower than 0.8 (but still satisfactory) were *Pallavicinia Lyiellii and Cephalozia macrostachya. Cephalozia macrostachya* is particularly problematic to identify and is thought to be under-represented in distribution maps. For this reason, although we have applied the target group sampling to correct for the sampling effort (see methods), a bias could still be present because the species might not have been detected in some surveys. *Cephalozia macrostachya* can in fact only be identified with certainty when fertile and needs to be identified at the right time of the year.

Another source of uncertainty is the coarse resolution of the model, which was necessary given the extension of the study area and the availability of climatic projections. This is likely to hide the effects of some local factors like micro-topography, that could have an impact on the real distribution of the species by creating different ecological niches (Lindsay et al., 2014).

Finally, something that needs to be noted is that also the current distribution of species is affected, besides by climate, by habitat loss that has happened during the last centuries because of different human activities. For this reason, the current modelled favourable climatic space of each species, which is based on recent observations (1980 -2018), might be wider than predicted: areas where the species are absent could in fact be climatically suitable for them, and their absence might be due to other factors like urban or agriculture development.

### 4.2 Present and future climatic suitability

Seven of the considered species are predicted to lose some extent of their climatic space. Four of them (*Sphagnum lindbergii, Sphagnum majus, Sphagnum riparium* and *Cephalozia loitlesbergeri)* have similar current distributions (limited to the north of the UK and to mountain areas in Northern Ireland, Wales and Devon), and are the species expected to face the greatest reduction in their environmental space (−100%, - 98%, – 90.5% and -53.9%). In Preston et al. (2013), these three mosses belong to the *Kiaeria falcata* cluster, which groups species located at high altitudes in Scotland, with mainly Arctic, Boreo-arctic and Boreal ranges, whose needs are acidic substrates, very cold winters and summers and very high rainfalls. *Cephalozia loitlesbergeri*, a liverwort, is included in the *Anastrepta orcadensis* cluster in Preston et al. (2011), with upland species that can be found in localities with low January and July temperature and very high precipitation. In our models, temperature-related variables give the greatest contribution to their distribution (see Appendices 4 and 5 in Supplementary material 1). It is not surprising – given their climatic needs – that they are predicted to lose such an extended area (with different intensities) mainly in the east and in the south of Scotland (except for *Sphagnum riparium* which maintains more areas in the east than in the west). Here, summer temperatures are already comparatively warm and are predicted to increase more than in the north and in the west.

Following the same classification (Preston et al., 2013), *Campylopus shawii* and *Sphagnum strictum* belong to the *Blindia acuta* cluster, which includes boreal and hyperoceanic upland species that need high rainfall and favour acidic, nutrient poor environments. According to our models, they have a similar distribution and they are both predicted to lose around one third of their current distribution in the 2050s (−34.1% and -33.8%), with a shift northwards and no suitable areas below the level of the Central Belt, probably due to a more marked increase in temperature (especially the maximum temperature of the warmest month, which, in the models of both of the species has a great relevance). They are also the species that mostly rely on winter precipitation, and this might explain the fact that, despite the overall loss of suitable area, they reinforce their presence in the west (especially *Campylopus shawii*), where an increase in precipitation, particularly during winter, is predicted. Something that we need to be cautious of, though, is that in spite of the predicted overall increase in rainfalls, there is still uncertainty with regard to its periodicity and to the possibility of drought periods, which might be more frequent/longer in the future and might be dangerous for species such as these that are intolerant to even short period of desiccation. The species that, overall, is losing the least favourable areas, is *Sphagnum phulchrum* (−4.4%). In Preston et al. (2013), this species belongs to the *Pleurozium schreberi* cluster, characterized by species with a wide geographic range, with a need for acidic, nutrient-poor and moist or wet substrates. Its current distribution is one of the most widespread in the UK among the considered species. Being already present in warmer areas (southern England), the predicted increase in temperature does not seem to affect its distribution in Scotland; in fact, our model shows an expansion in the south. Species with a similar, although more accentuated, distribution are *Cephalozia macrostachya* and *Pallavicinia lyellii*, which are also the only species that according to our models will increase their area. In Preston et al. (2011) they are included in the *Cladopodiella fluitans* cluster, with species distributed in areas with warmer than average temperatures and lower rainfalls, which could explain this expansion, especially towards the east.

### 4.3 Considering blanket bog projections

The blanket bog layer was included as an explanatory variable in the models of seven species. When projected change of blanket bog was included, the predicted distribution of all species was almost always different from the distributions obtained keeping the variable constant. We used two blanket bog models, BT and LM, equally plausible but based on different combination of variables (see Clark et al., 2010 and Ferretto et al., 2019), that show the blanket bog not at risk of loss in the 2050s according to the medium emission scenario (fig. 2). In general, changes of predicted species distribution were more substantial when we used BT rather than LM. The only species whose distribution remains substantially the same if we account for the change in blanket bog habitat are *Campylopus shawii* and *Sphagnum strictum* and this makes intuitive sense because their distribution is already almost entirely confined to areas where blanket bog is not predicted to be at risk. Despite the similar outcome for these two species, in the case of *Sphagnum strictum*, the blanket bog variable is not relevant for the model, while it is very important for *Campylopus shawii* (see Appendices 4 and 5 in Supplementary material 1). The non-relevance of the blanket bog layer for *Sphagnum strictum* is also confirmed by the fact that it is not an obligated blanket bog species, but it is usually found in wet heat, associated with *Molinia*, in boggy grasslands and in thin blanket bogs (Blockeel et al., 2014). Blanket bog has shown to be among the most important variables for *Pallavicinia lyellii* and for *Cephalozia macrostachya* and for these species it is evident that there is a predicted retreat in the areas where blanket bog is predicted to be at risk. While *Cephalozia macrostachya* is a typical blanket bog species, *Pallavicinia lyellii* grows on bog margins and in woodlands. The importance attributed by the model to the blanket bog layer for *Pallavicinia lyellii* might be due to the spatial uncertainty of the observed records, which is up to 1 km, so that a record classified as “on blanket bog” might have been found on bog margins instead.

The fact that for some species there is a substantial difference when the changes in the habitat are considered indicates the importance of its inclusion in the models for those species that have a strong link with blanket bogs. This obviously has implications for previous species distribution modelling work. For example, Smart et al. (2010) modelled the distribution of ombrotrophic *Sphagnum* species in the UK and projected it in the 2050s considering changes in climatic conditions and level of pollutants. According to their model, areas in the Northwest of Scotland will be the most negatively impacted, whereas in the East, the impact will be weaker. It would be interesting to add the changes in blanket bog distribution predicted by bioclimatic models to see if this changes their predictions.

An alternative approach, that would allow refinement of bioclimatic models to account for habitat, could be to include in the SDMs only environmental variables and to use habitat distribution as a mask on the model outputs. Stanton et al. (2012) compared the two methods and found that using habitat distribution as a mask on SDMs predictions returned poorer performances than including habitat distribution among the variables. In our study, blanket bog is the main habitat for the considered species, but not all the records were found on blanket bogs, and applying the mask would have excluded all the areas where blanket bogs are at risk but that could still support the species.

### 4.4 Implications for conservation and peatland restoration

SDMs can give useful information for conservation purposes. In spite of their limitations, when used at a coarse scale they represent a quick and valuable input for planning conservation measures. They have received criticism because they do not account for many factors (Sinclair et al., 2010). Among these, there are dispersion, adaptation, feedbacks and habitat availability. These are all important elements, but while climate data are widely available, data about interactions, adaptation and other more specific mechanisms are more difficult to obtain for modelling large areas, and their level of uncertainty is high.

In light of this, and while acknowledging their limitations, SDMs can be very useful as a first approach for modelling species distribution changes over large areas (for example for an entire country) and the inclusion of habitat is an easy addition given that land cover maps, habitat maps and satellite data are now readily available. Such an addition is likely to be particularly important when the species has a strong relationship with a specific habitat, and when habitat change can thus change substantially the modelled distribution, bringing it one step closer to the realized niche. But, in order for SDMs to be informative for future projections we cannot assume that the habitat will always be constant, especially in light of climate change and land use change, that are threatening many habitats (Mantyka-Pringle et al., 2012). If future projections of habitat are not directly available, the IUCN criteria could be used for assessing the risks to ecosystems (Bland et al., 2017) and they could be a start in defining the future distribution of habitats. In the specific case of blanket bog in Scotland, given that their importance for climate change mitigation has now been acknowledged and there are plans for their protection from other direct anthropogenic threats like peat extraction, afforestation and drainage for agriculture, the main threat to their survival is linked to the draining effects of climate change. For this reason, using only bioclimatic models for blanket bog projections is reasonable. In other cases, considering a combination of climatic threats and direct human pressures might be necessary. For example, tropical peatlands are threatened also by land use change, e.g. to leave space for palm oil plantations (Wicke et al., 2011) that might have a higher impact than climate change on this habitat.

In the specific case of rare species, this first evaluation of future species distribution could help the planning of protected site networks, allowing areas that are predicted not to be suitable for the species in the future to be avoided, or to focus more efforts in areas that are not predicted to be at risk. Considering the simplifications of this method and the uncertainties related to the omission of important ecological factors, a further and more accurate analysis should then be conducted in the selected areas, but including the habitat change in the first step can bring an important improvement without increasing the effort and the costs of the evaluation.

This approach could also be included in the assessment of the IUCN categories in the Red List of species. In the Red List, the status of a species is evaluated through different criteria, based on “observed, estimated, inferred or projected” data (IUCN, 2019). The use of SDMs is already included in the IUCN guidelines, providing that it is regulated and that the output is carefully interpreted and integrated with other important ecological factors like, for example, dispersal capability and physiological characteristics. The inclusion of land use as a variable of the model is already suggested in order to avoid an overestimation of the suitable areas (IUCN, 2019). One of the criteria for which a species can be included in the Red List is a continuing decline in its geographical range, in the form of Extent of Occurrence (EOO) and/or Area of Occupancy (AOO) (IUCN, 2019). Including habitat change in SDMs can improve the assessment of the trends of habitat suitability for a species, which can then be used to help identify any declines in EOO and AOO.

Knowing where species are more or less vulnerable can also help planning the peatland restoration activities that are now being carried out in many countries in all the continents (Cris et al., 2014). Peatlands are in fact estimated to currently store over 600 Gt of carbon, 547 Gt of which is in northern peatlands (Yu et al., 2010), a great amount of which could be released into the atmosphere due the degradation of wide areas of this habitat. Restoration projects aim to avoid this and to ensure that degraded areas will return to an “active” state (i.e. to sequester carbon), and the role of vegetation is essential to rebuild the mechanisms that will bring back this ecosystem function. For example, in the specific case of Scotland, applying this approach to important habitat forming species and more common species like *Sphagnum capillifolium* and *Sphagnum papillosum*, which are all keystone species present in Scottish blanket bogs, could inform revegetation activities in peatland restoration projects. Being common species, the number of observations would also be substantial compared to the number of records used in this study, returning more robust results. Understanding how habitat and species distribution are affected by climate change and focussing on their interconnections is important in order to plan peatland restoration in an effective way bringing benefits both to conservation (in the case of rare species) and to carbon sequestration strategies (in the case of more common species with a high value in these terms). In our specific case, the threats for blanket bog that seem to have the greatest extra-impact on most of the considered species are in the North of Scotland in the Flow Country, where most of the restoration efforts are focussed, in part because of the substantial amounts of carbon stored in this area (Hermans et al., 2019) but most of all because of the positive effect of forest to bog restoration for biodiversity (Wilson et al., 2014). If we had not included the predicted distribution of blanket bogs, this area would have appeared suitable in the future for most of the species that we have analysed. However, this is not true for two of the selected species, where blanket bog was excluded by the variable selection process, and in another species, where although it was included, it did not increase the gain of the model. The inclusion of the habitat can hence be fundamental for some species but not crucial for others, and this should be explored beforehand, because omitting the change of habitat *a priori* can overestimate the potential projections in some areas if the species are strictly linked to the habitat and the habitat is predicted to face unsuitable climatic conditions. In the context of restoration, our approach could also be helpful if applied to other more common species that are considered in revegetation programs, both in the choice of the species that might be more adaptable, and in the choice of where to address most of the efforts to apply these kind of interventions.

## 5. Conclusion

Although SDMs represent a first approximation of the genuine distribution of a species, given the availability of climate data and climate projections they can also return quick and valuable information at a coarse scale about potential future distributions. However, these future projections might only represent a rough approximation of the future distribution of the species because they do not include many other important factors. To this end, considering the future distribution of habitat alongside climatic variables should make them more realistic, but if SDMs are projected into the future, the assumption that the habitat will remain constant is questionable. We tested the impact of future projections of habitat change on the distributions predicted by SDMs. Our results showed that the predicted future distribution can be quite different if habitat changes are accounted for, particularly when there is a strong relationship between the species and its habitat. This simple addition can improve the reliability of SDMs in the first steps of planning for conservation and restoration.

## Notes

### Competing Interest Statement

The authors have declared no competing interest.

