## Supplementary material for "Modelling the future distribution of rare bryophytes in Scotland: is inclusion of habitat loss important?"

### Appendix 1

*Table S1.1: Sources of species data. Data Provider, Original Recorder [where identified], and the NBN Trust bear no responsibility for any further analysis or interpretation of that material, data and/or information.*

| Name | doi | Licence |
|---|---|---|
| BIS for Powys & Brecon Beacons National Park | | |
| BRERC species records from all years at full resolution excluding Notable Species within the last 10 years | doi:10.15468/h1ln5p | Creative Commons, with Attribution, Non-commercial v4.0 (CC-BY-NC) BRERC retains copyright. The data is provided via the NBN Atlas for non-commercial purposes. |
| Bristol Regional Environmental Records Centre | | |
| British Bryological Society | | |
| Bryophyte data for Great Britain and Ireland from the British Bryological Society held by BRC: data compiled post-Atlas | doi:10.15468/ttzehy | Creative Commons with Attribution 4.0 (CC-BY) CC-BY |
| Bryophyte data for Great Britain from the British Bryological Society held by BRC: Atlas 2014 | doi:10.15468/gvqhjb | Creative Commons with Attribution 4.0 (CC-BY) CC-BY |
| Bryophyte records via iRecord | 10.15468/2g3nyq | Creative Commons with Attribution 4.0 (CC-BY) CC BY |
| Bryophyte Survey of the Poole Basin Mires - NBN South West Pilot Project Case Studies | doi:10.15468/eklhxs | Creative Commons, with Attribution, Non-commercial v4.0 (CC-BY-NC) CC-BY-NC |
| Centre for Environmental Data and Recording | | |
| Commissioned surveys and staff surveys and reports for Scottish Wildlife Trust reserves - Verified data | doi:10.15468/a6snhl | Creative Commons with Attribution 4.0 (CC-BY) CC-BY |
| Cumbria Biodiversity Data Centre | | |
| Data from Defra Family Organisations supplied to Staffordshire Ecological Record | doi:10.15468/giebpp | Creative Commons, with Attribution, Non-commercial v4.0 (CC-BY-NC) CC-BY-NC |
| Dorset Environmental Records Centre | | |
| Dorset Sites of Nature Conservation Interest (SNCI) species records 2000-2008 | doi:10.15468/r1ebqb | Creative Commons, with Attribution, Non-commercial v4.0 (CC-BY-NC) CC-BY-NC |
| Dorset Sites of Nature Conservation Interest (SNCI) species records pre 2000 | doi:10.15468/qyg29v | Creative Commons, with Attribution, Non-commercial v4.0 (CC-BY-NC) CC-BY-NC |
| Dorset SSSI Species Records 1952 - 2004 (Natural England) | doi:10.15468/vcjzts | Creative Commons, with Attribution, Non-commercial v4.0 (CC-BY-NC) CC-BY-NC |
| HBRG Fungus, Lichen & Lower Plants Dataset | doi:10.15468/rt48my | Creative Commons with Attribution 4.0 (CC-BY) CC-BY |
| Highland Biological Recording Group | | |
| John Muir Trust | | |
| Kent & Medway Biological Records Centre | | |
| Lancashire Environment Record Network | | |
| LERN Records | doi:10.15468/esxc9a | Creative Commons, with Attribution, Non-commercial v4.0 (CC-BY-NC) CC-BY-NC |
| Miscellaneous records held by BIS | doi:10.15468/mo7peo | Creative Commons, with Attribution, Non-commercial v4.0 (CC-BY-NC) |
| Montgomeryshire Wildlife Trust records held by BIS | doi:10.15468/vozyfp | Creative Commons, with Attribution, Non-commercial v4.0 (CC-BY-NC) |

| | | |
|---|---|---|
| Mosses and Liverworts: Records for Kent. | doi:10.15468/wlgnsv | Creative Commons, with Attribution, Non-commercial v4.0 (CC-BY-NC) CC-BY-NC |
| National Trust for Scotland Species Records | doi:10.15468/a5y1cz | Creative Commons with Attribution 4.0 (CC-BY) CC-BY |
| National Trust Species Records | doi:10.15468/opc6g1 | Creative Commons, with Attribution, Non-commercial v4.0 (CC-BY-NC) CC-BY-NC |
| Natural Resources Wales Regional Data : Mid-Wales | doi:10.15468/whj6d7 | Open Government Licence (OGL) OGL |
| NatureScot | | |
| North East Scotland Biological Records Centre | | |
| North East Scotland Mosses and Liverworts (1950-2014) | doi:10.15468/wgzruc | Creative Commons, with Attribution, Non-commercial v4.0 (CC-BY-NC) CC-BY-NC |
| Northern Ireland Environment Agency (NIEA) Collated Species Records | | Open Government Licence (OGL) |
| NRW Regional Data: all taxa (excluding sensitive species), West Wales | doi:10.15468/q3d1hl | Open Government Licence (OGL) OGL |
| Phase 2 Lowland Peatland Survey of Wales 2004 onwards | doi:10.15468/ijegqu | Open Government Licence (OGL) OGL |
| Plants, Bryophytes and Lichens recorded on the Nevis Estate during summer 2003. | doi:10.15468/vgoj2i | Creative Commons, with Attribution, Non-commercial v4.0 (CC-BY-NC) CC-BY-NC |
| RECORD Bryopsida Data | doi:10.15468/coah7z | Creative Commons, with Attribution, Non-commercial v4.0 (CC-BY-NC) CC-BY-NC |
| Royal Botanic Garden Edinburgh Herbarium (E) | | Creative Commons, with Attribution, Non-commercial v4.0 (CC-BY-NC) Copyright Royal Botanic Garden Edinburgh. Contact us for rights to commercial use. |
| Scottish Wildlife Trust | | |
| SER Site-based Surveys | doi:10.15468/h2yko0 | Creative Commons, with Attribution, Non-commercial v4.0 (CC-BY-NC) CC-BY-NC |
| Shropshire Ecological Data Network database | doi:10.15468/5v5pvk | Creative Commons with Attribution 4.0 (CC-BY) CC-BY |
| Staffordshire Ecological Record | | |
| Standing Waters Database - Scotland | doi:10.15468/iwllx8 | Open Government Licence (OGL) OGL |
| Suffolk Biodiversity Information Service (SBIS) Dataset | doi:10.15468/ab4vwo | Creative Commons, with Attribution, Non-commercial v4.0 (CC-BY-NC) CC-BY-NC |
| Tullie House Museum Natural History Collections | doi:10.15468/epewfs | Creative Commons, with Attribution, Non-commercial v4.0 (CC-BY-NC) CC-BY-NC |
| Vascular Plants and Bryophytes of Glen Sligachan | doi:10.15468/e8s4tr | Creative Commons, with Attribution, Non-commercial v4.0 (CC-BY-NC) CC-BY-NC |
| West Wales Biodiversity Information Centre | | |
| WTSWW Data: All Taxa (West Wales) | doi:10.15468/gaakk2 | Creative Commons, with Attribution, Non-commercial v4.0 (CC-BY-NC) CC-BY-NC |

### Appendix 2

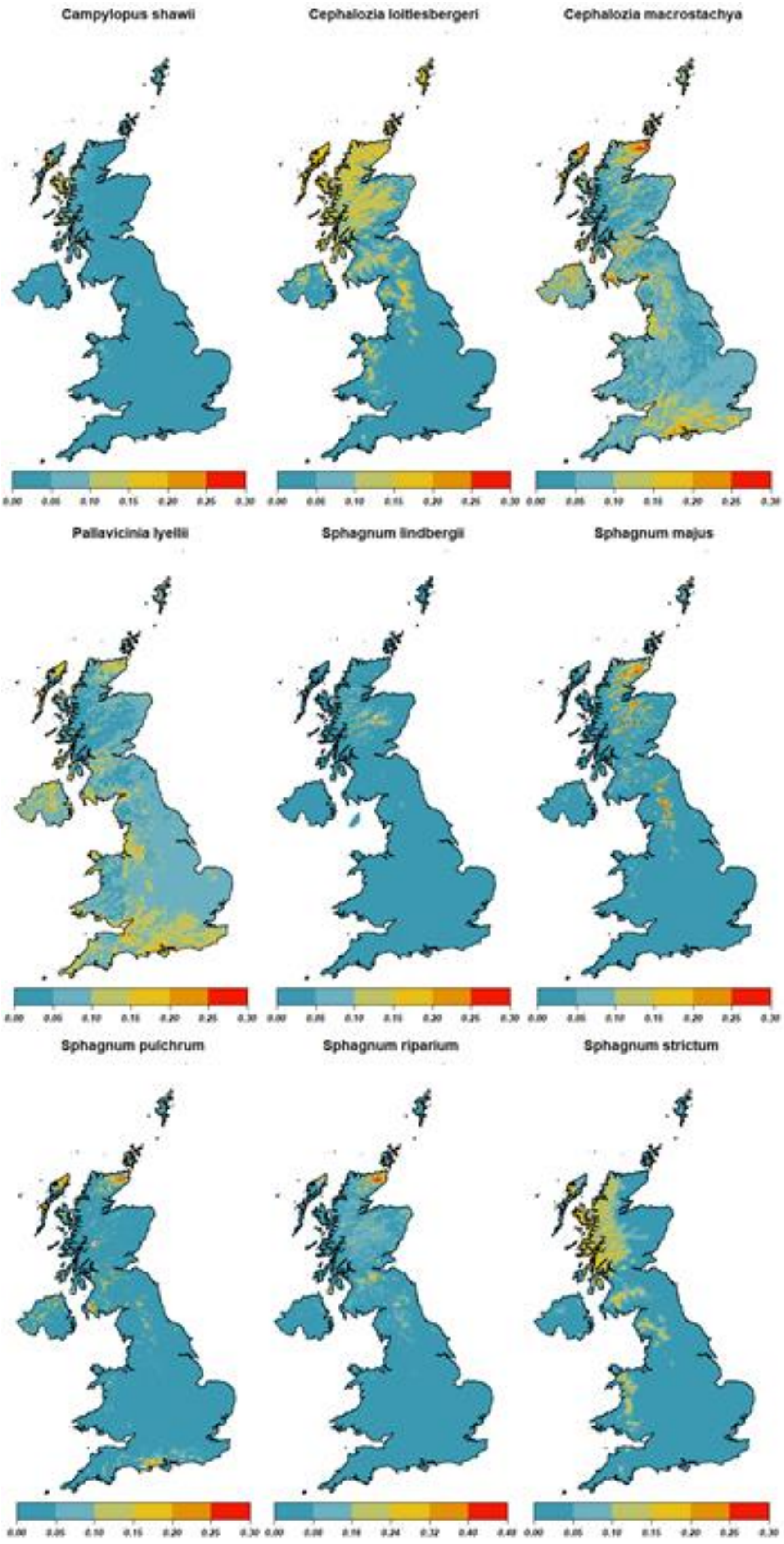


*Figure S2.1 Maxent probability distribution for all the studied species of bryophytes.*

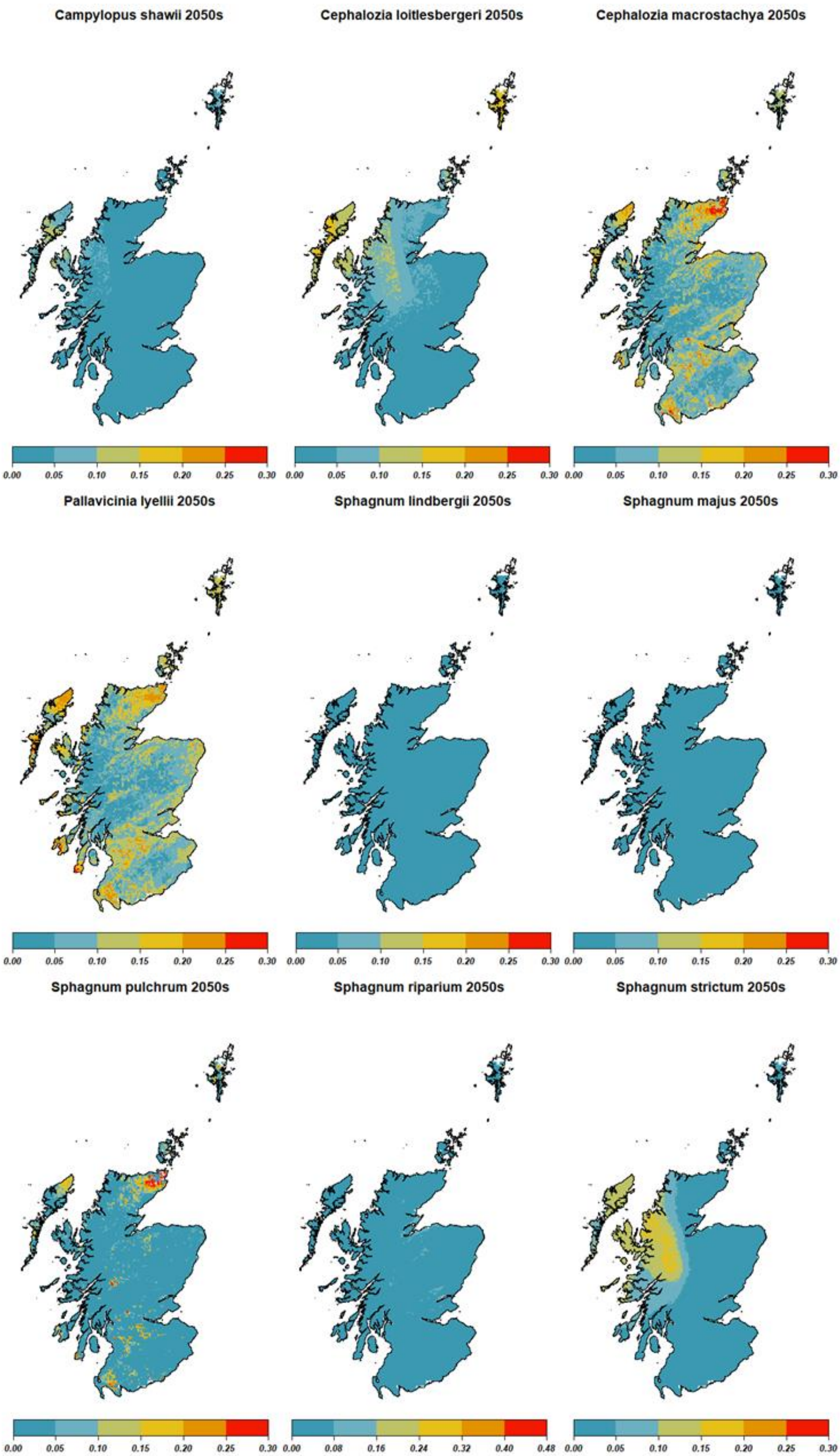


*Figure S2.2 Maxent future probability distribution for all the studied species of bryophytes, obtained changing only the climate variables.*

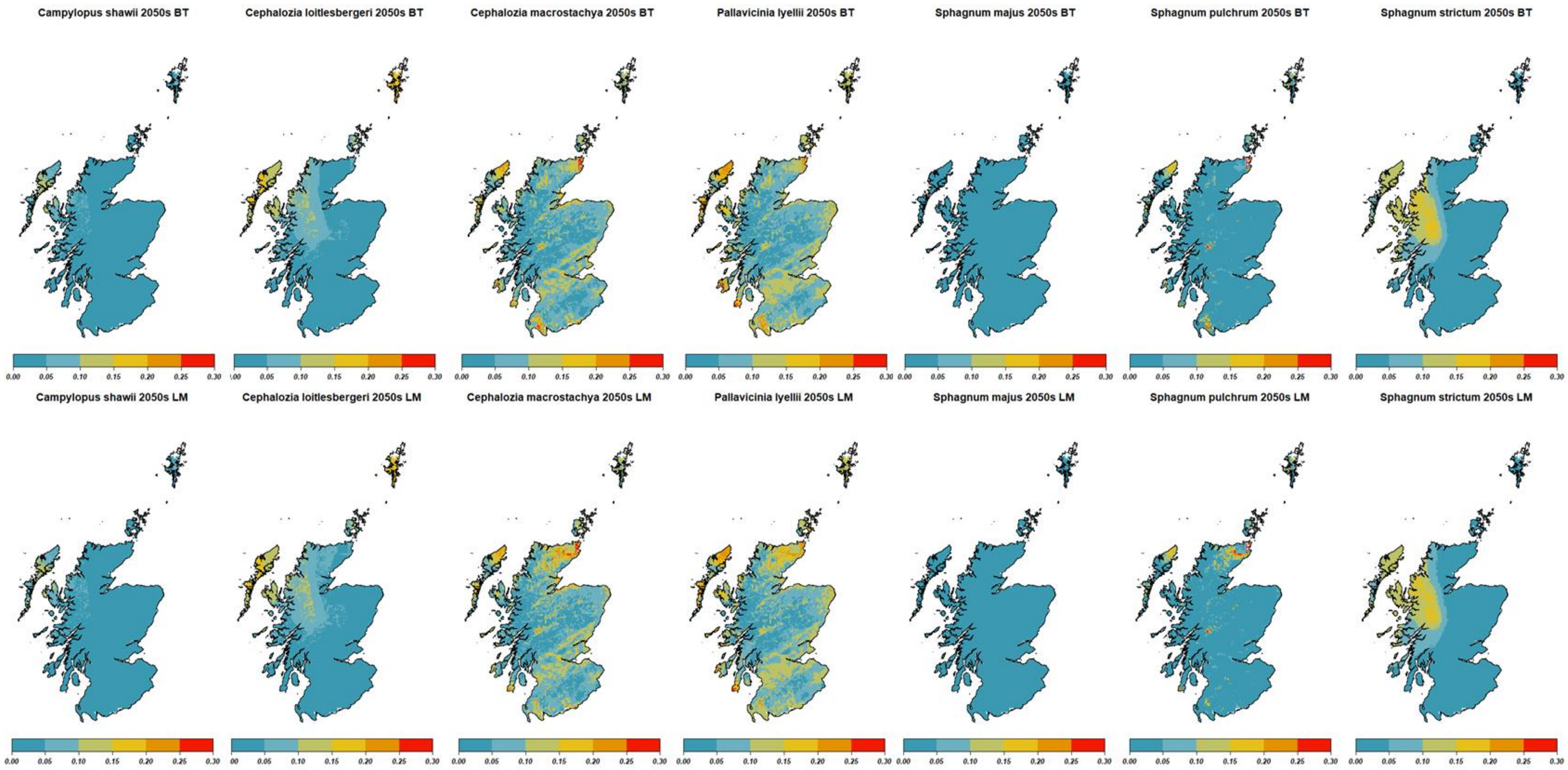


*Figure S2.3 Maxent future probability distribution for all the studied species of bryophytes using BT and LM modelled blanket bog distribution*

### Appendix 3

The equations of BBOG TREE model (1) and LM model (2) used in Ferretto et al. (2019) are the following:

BBOG TREE:

Probability of Blanket Peat = 1 if $T_{max} < 17.4^{o}C$ and TMI > 0.41

or

$T_{max} > 17.4^{o}C$ *and* AAMWD < -28.6 mm (1)

where

$T_{max}$ = maximum yearly temperature

TMI = Thornthwaite – Mather Moisture Index (which is a measure of the annual balance between precipitation and potential evapotranspiration)

AAMWD = annual accumulated monthly water deficit (which, in contrast to TMI, accounts for the seasonality in the balance between precipitation and potential evapotranspiration).

LM:

Probability of Blanket Peat = 1 if P > 1000 mm and $T_m < 15^{o}C$ (2)

where

P = total yearly precipitation

Tm = maximum monthly mean temperature

### Appendix 4

*Table S4.1: Contribution (in percentage) of each variable to model each species distribution. NA values mean that the variable does not enter the model (it was excluded a priori through the variable selection process).*

| | *Campylopus shawii* | *Cephalozia loitlesbergeri* | *Cephalozia machrostachya* | *Pallavicinia lyellii* | *Sphagnum lindbergii* | *Sphagnum majus* | *Sphagnum pulchrum* | *Sphagnum riparium* | *Sphagnum strictum* |
|---|---|---|---|---|---|---|---|---|---|
| Blanket bog | 20.0 | 12.1 | 31.7 | 22.9 | NA | 45.1 | 15.0 | NA | 0.2 |
| Slope | 1.8 | 0 | 32.6 | 42.1 | NA | 10 | 33.6 | 12.4 | 0 |
| Mean t˚ | 0 | 2.9 | NA | 2.2 | 98.4 | 44.9 | 0.2 | 78.1 | 0.4 |
| Total precipitation | 0 | NA | NA | NA | NA | NA | 1.9 | 0.3 | NA |
| Max t˚ of the warmest month | 51.5 | 84.9 | 4.9 | 0.1 | NA | NA | 0.2 | 3.9 | 44.6 |
| Min t˚ of the coldest month | 7.6 | NA | 0.8 | NA | 0.2 | NA | 1.5 | 0.3 | NA |
| Summer precipitation | 0 | NA | 0.9 | NA | NA | NA | 15.3 | 0 | 0.1 |
| Winter precipitation | 19.0 | 0.1 | 29.1 | 9.4 | NA | 0 | 32.3 | 4.9 | 54.4 |
| PET | NA | NA | NA | 3.6 | 1.5 | NA | NA | NA | 0.3 |

### Appendix 5

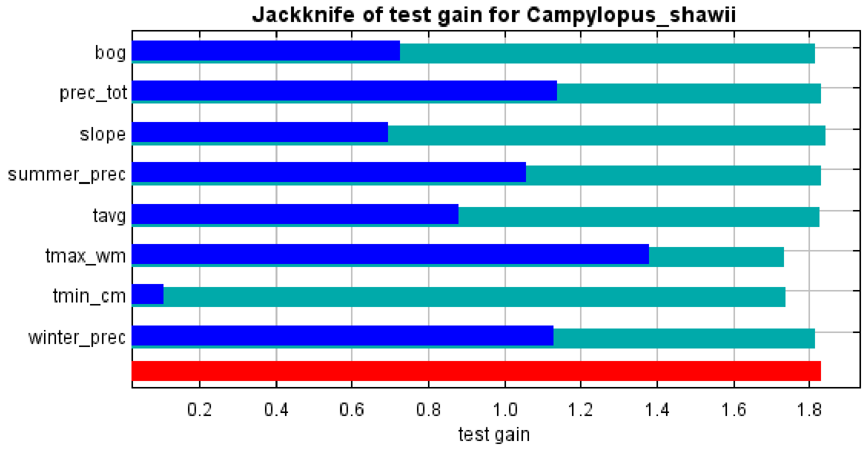
Jackknife of test gain for Campylopus_shawii
bog
prec_tot
slope
summer_prec
tavg
tmax_wm
tmin_cm
winter_prec
0.2
0.4
0.6
0.8
1.0
1.2
1.4
1.6
1.8
test gain

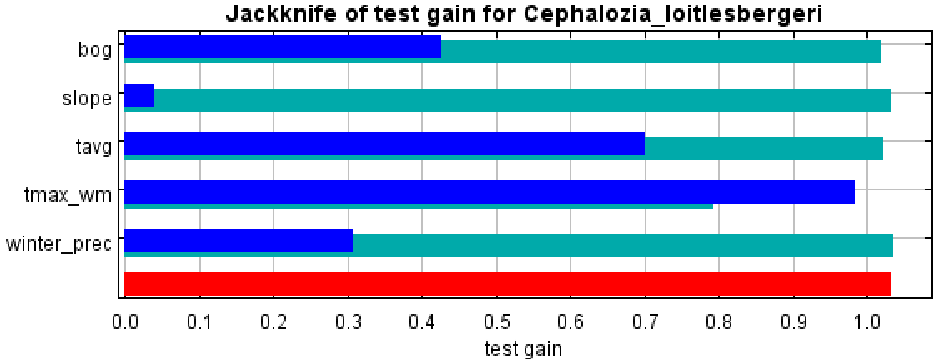
Jackknife of test gain for Cephalozia_loitlesbergeri
bog
slope
tavg
tmax_wm
winter_prec
0.0
0.1
0.2
0.3
0.4
0.5
0.6
0.7
0.8
0.9
1.0
test gain

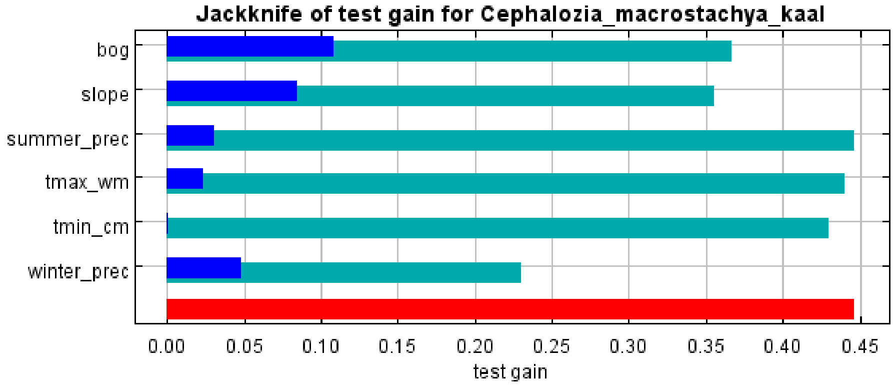
Jackknife of test gain for Cephalozia_macrostachya_kaal
bog
slope
summer_prec
tmax_wm
tmin_cm
winter_prec
0.00
0.05
0.10
0.15
0.20
0.25
0.30
0.35
0.40
0.45
test gain

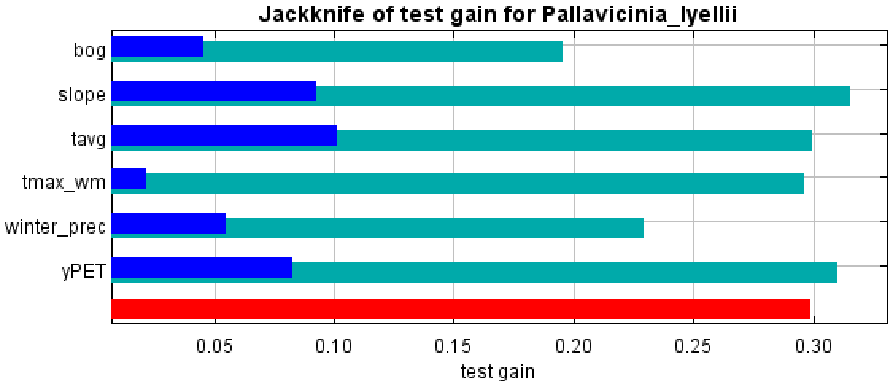
Jackknife of test gain for Pallavicinia_lyellii
bog
slope
tavg
tmax_wm
winter_prec
yPET
0.05
0.10
0.15
0.20
0.25
0.30
test gain

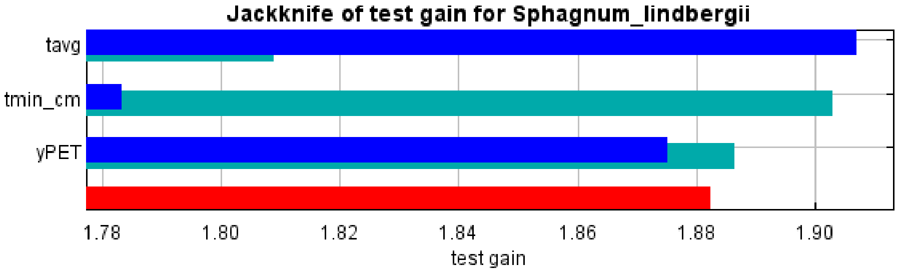
Jackknife of test gain for Sphagnum_lindbergii
tavg
tmin_cm
yPET
1.78
1.80
1.82
1.84
1.86
1.88
1.90
test gain

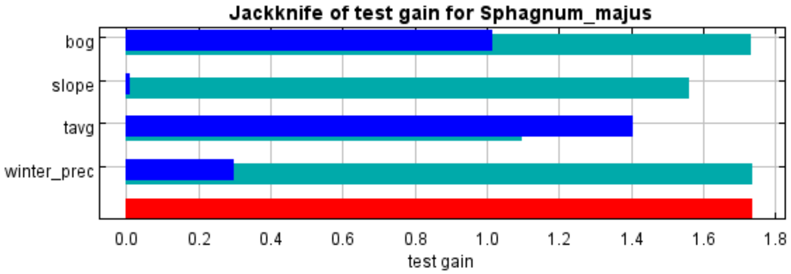
Jackknife of test gain for Sphagnum_majus
bog
slope
tavg
winter_prec
0.0
0.2
0.4
0.6
0.8
1.0
1.2
1.4
1.6
1.8
test gain

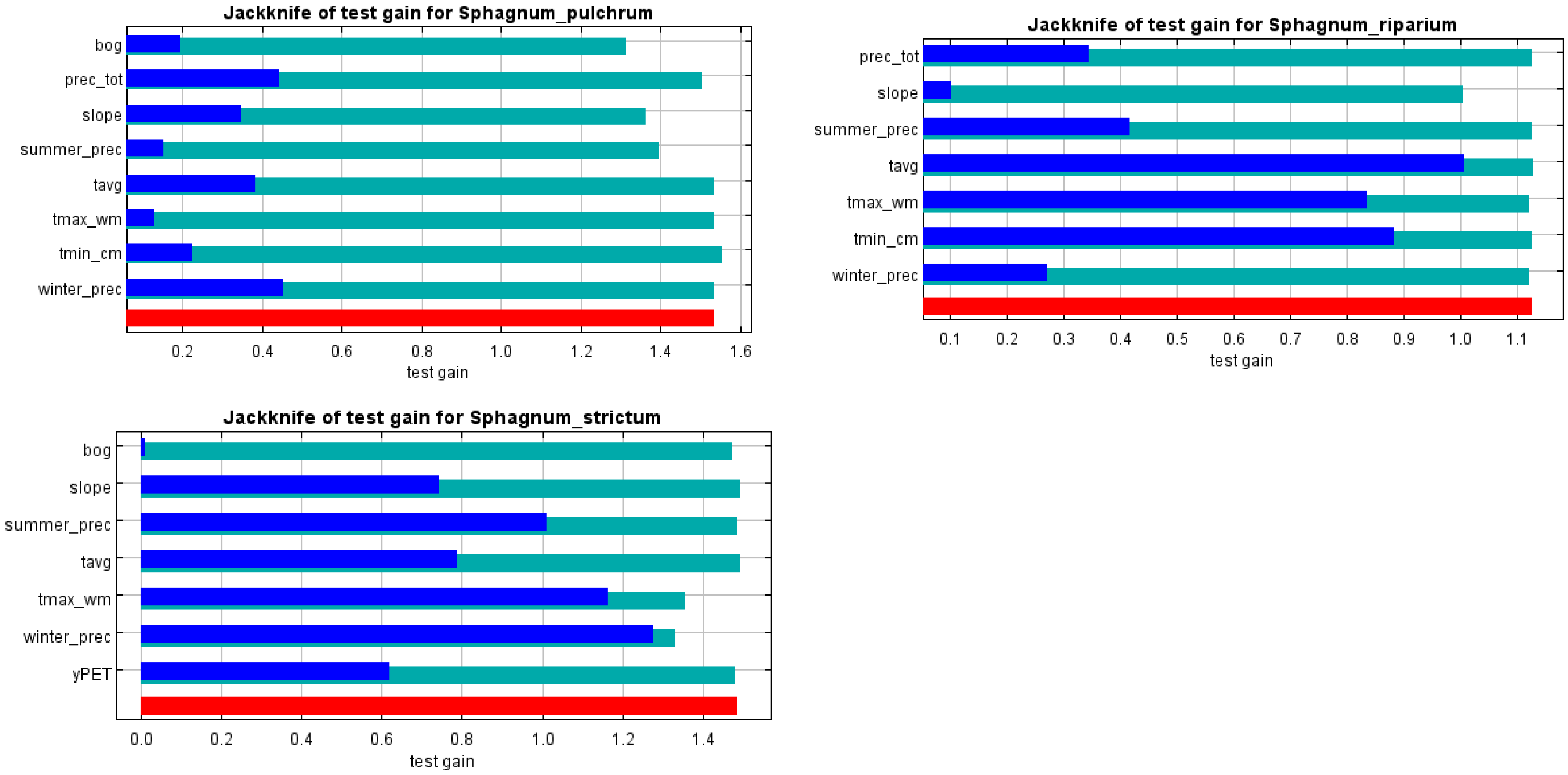


*Figure S5.1: Outputs of the jackknife test produced by Maxent for each species. In light blue: test gain obtained by excluding the correspondent variable. In dark blue: test gain obtained by using only the correspondent variable. In red: test gain obtained including all the variables.*

### Appendix 6

In this appendix we report the results of preliminary analysis that include also pollution data ($SO_x$, $NO_x$ and $NH_x$ deposition), that were not included in the main document because, being at different resolution, they increase uncertainty.

For the UK, cumulative deposition data for NHx (kg N $ha^{-1}$ $yr^{-1}$), NOx (kg N $ha^{-1}$ $yr^{-1}$) and SOx (kg S $ha^{-1}$ $yr^{-1}$) were obtained at 5 km resolution from the CBED model (Smith et al., 2000) adopting historical scaling factors (Fowler et al., 2004) on a base year of 2005, and averaged for the period 2004-2006. SOx deposition has greatly decreased in the UK from the late 1980s (RoTAP, 2012), but since the number of records in the 1980s is very small compared to the overall number of records, we think that this change is not going to affect the results.

We assumed that in the 2050s, NHx, NOx and SOx deposition is not going to change greatly from the current values, so from the CBED model we selected the average for the period 2017-2019.

The model was run following the methods reported in the main text, with the only difference that pollution data were also included in the process.

The following tables report the model performances for all the species, the combination of variables and β values that minimize the AIC score, the contribution of each variable, the thresholds used to convert the probability maps into presence/absence maps and the % of changes predicted by the model in the future.

*Table S6.1 Test AUC values and their standard deviation and Boyce Index for each species.*

| Species | Average test AUC | Standard deviation | Boyce Index |
|---|---|---|---|
| *Campylopus shawii* | 0.947 | 0.015 | 0.899 |
| *Cephalozia loitlesbergeri* | 0.900 | 0.010 | 0.655 |
| *Cephalozia macrostachya* | 0.794 | 0.058 | 0.916 |
| *Pallavicinia lyellii* | 0.627 | 0.107 | 0.796 |
| *Sphagnum lindbergii* | 0.958 | 0.014 | 0.954 |
| *Sphagnum majus* | 0.947 | 0.013 | 0.671 |
| *Sphagnum pulchrum* | 0.925 | 0.020 | 0.779 |
| *Sphagnum riparium* | 0.888 | 0.022 | 0.935 |
| *Sphagnum strictum* | 0.921 | 0.014 | 0.878 |

Given the low AUC value for *Pallavicinia lyellii*, this species was excluded from the analysis.

*Table S6.2 Regularization multiplier (β) values and variables used to run Maxent.*

| | *Campylopus shawii* | *Cephalozia loitlesbergeri* | *Cephalozia macrostachya* | *Sphagnum lindbergii* | *Sphagnum majus* | *Sphagnum pulchrum* | *Sphagnum riparium* | *Sphagnum strictum* |
|---|---|---|---|---|---|---|---|---|
| β value | 4 | 5 | 4.5 | 3 | 3.5 | 2 | 4 | 4.5 |
| Blanket bog | ● | ● | ● | | ● | ● | | |
| Slope | | | ● | | ● | ● | | ● |
| Mean t˚ | | ● | | ● | ● | ● | ● | ● |
| Total precipitation | ● | | ● | | | ● | | |
| Max t˚ of the warmest month | | ● | ● | ● | | ● | | ● |
| Min t˚ of the coldest month | | | | ● | ● | ● | ● | |
| Summer precipitation | ● | | ● | | | ● | | |
| Winter precipitation | ● | | ● | | | ● | ● | ● |
| PET | | | | ● | | ● | | |
| NOx | ● | ● | | | | ● | ● | ● |
| NHx | ● | ● | | ● | ● | ● | ● | ● |
| SOx | | | ● | | | ● | ● | |

*Table S6.3: Contribution (in percentage) of each variable to model each species distribution. NA values mean that the variable does not enter the model (it was excluded a priori through the variable selection process).*

| | *Campylopus shawii* | *Cephalozia loitlesbergeri* | *Cephalozia macrostachya* | *Sphagnum lindbergii* | *Sphagnum majus* | *Sphagnum pulchrum* | *Sphagnum riparium* | *Sphagnum strictum* |
|---|---|---|---|---|---|---|---|---|
| Blanket bog | 5.2 | 12.7 | 53.0 | NA | 62.9 | 8.7 | NA | NA |
| Slope | NA | NA | 27.6 | NA | 0.1 | 37.7 | NA | 0.5 |
| Mean t° | NA | 0 | NA | 70.8 | 33.7 | 1.4 | 92.8 | 1.4 |
| Total precipitation | 17.9 | NA | 0 | NA | NA | 0.6 | NA | NA |
| Max t° of the warmest month | NA | 73.5 | 0 | 15.2 | NA | 0 | NA | 12.7 |
| Min t° of the coldest month | NA | NA | NA | 9.7 | 0.2 | 1.5 | 0 | NA |
| Summer precipitation | 0 | NA | 0.1 | NA | NA | 11 | NA | NA |
| Winter precipitation | 13.1 | NA | 18.8 | NA | NA | 34.7 | 2 | 60.1 |
| PET | NA | NA | NA | 3.4 | NA | 1 | NA | NA |
| $SO_x$ | NA | NA | 0.5 | NA | NA | 0.2 | 0 | NA |
| $NO_x$ | 57.1 | 0.2 | NA | NA | NA | 3 | 0.9 | 0 |
| $NH_x$ | 6.7 | 13.7 | NA | 0.9 | 3.2 | 0.2 | 4.2 | 25.3 |

*Table S6.4 Maxent thresholds for the presence/absence conversion, obtained maximizing the sum of sensitivity and specificity in the training dataset*

| Species | Presence/absence threshold |
|---|---|
| *Campylopus shawii* | 0.0366 |
| *Cephalozia loitlesbergeri* | 0.0632 |
| *Cephalozia macrostachya* | 0.1068 |
| *Sphagnum lindbergii* | 0.0296 |
| *Sphagnum majus* | 0.0204 |
| *Sphagnum pulchrum* | 0.0440 |
| *Sphagnum riparium* | 0.1105 |
| *Sphagnum strictum* | 0.0520 |

*Table S6.5 Change in the predicted future distribution (2050s) compared to the current modelled distribution for each species, obtained considering only climate variables or climate variables and predicted blanket bog distribution according to BT and LM models*

| *Species* | *% of change* | *% of change BT bog distribution* | *% of change LM bog distribution* |
|---|---|---|---|
| *Campylopus shawii* | *+50.1* | *+47.8* | *+49.6* |
| *Cephalozia loitlesbergeri* | *-44.8* | *-48.8* | *-46.8* |
| *Cephalozia macrostachya* | *+16.6* | *-5.59* | *+6.06* |
| *Sphagnum lindbergii* | *-100* | */* | */* |
| *Sphagnum majus* | *-63* | *-81.7* | *-68.72* |
| *Sphagnum pulchrum* | *-20* | *-43.1* | *-34.5* |
| *Sphagnum riparium* | *-94.8* | */* | */* |
| *Sphagnum strictum* | *+24.5* | */* | */* |

The following maps show the results of the model run in the UK in the present and in Scotland in the 2050s according to the medium emission scenario. Future projections are shown both without changing the blanket bog layer and changing the blanket bog layer according to the BT and LM blanket bog distribution.

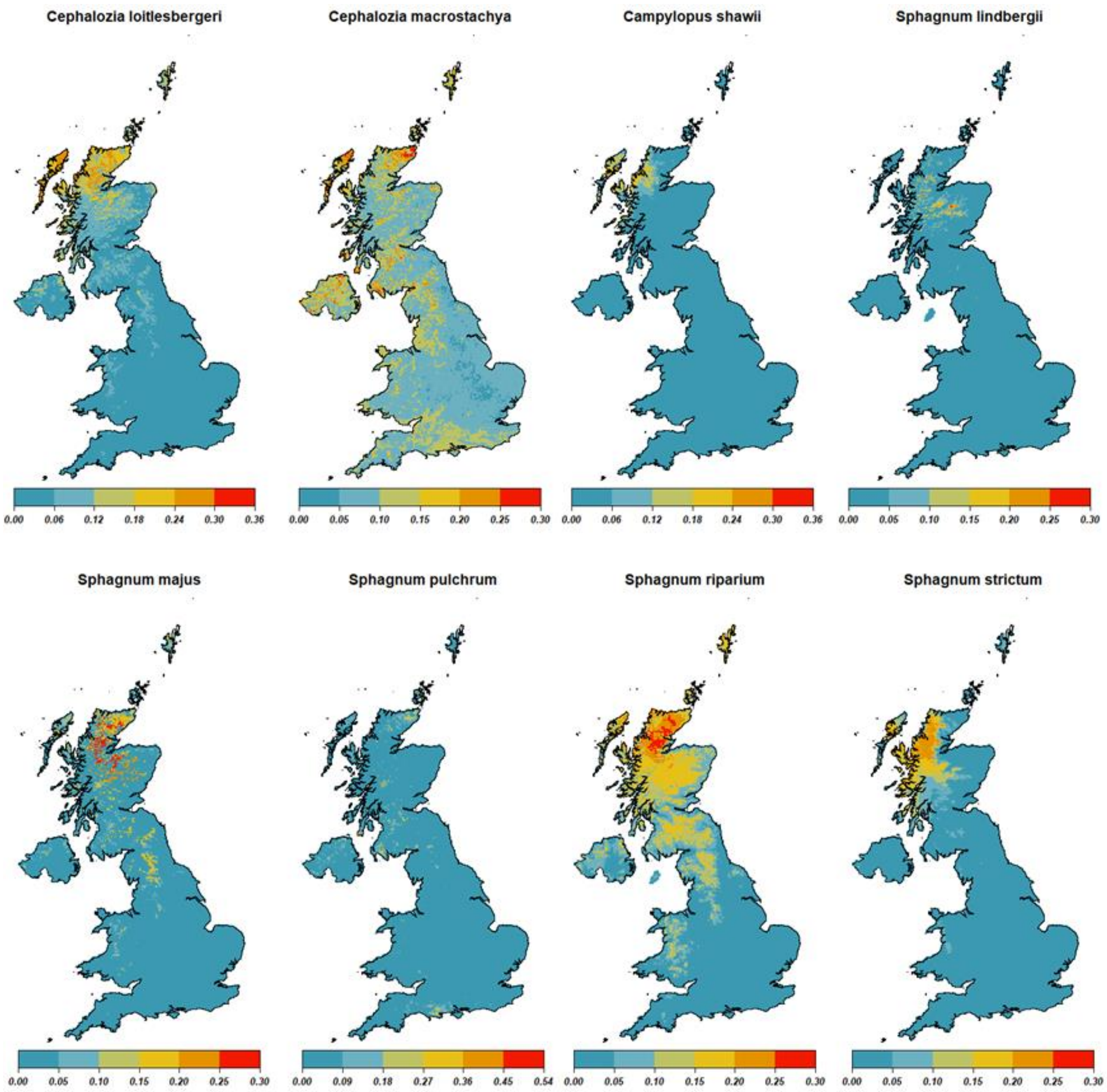


*Figure S6.1 Maxent probability distribution for all the studied species of bryophytes.*

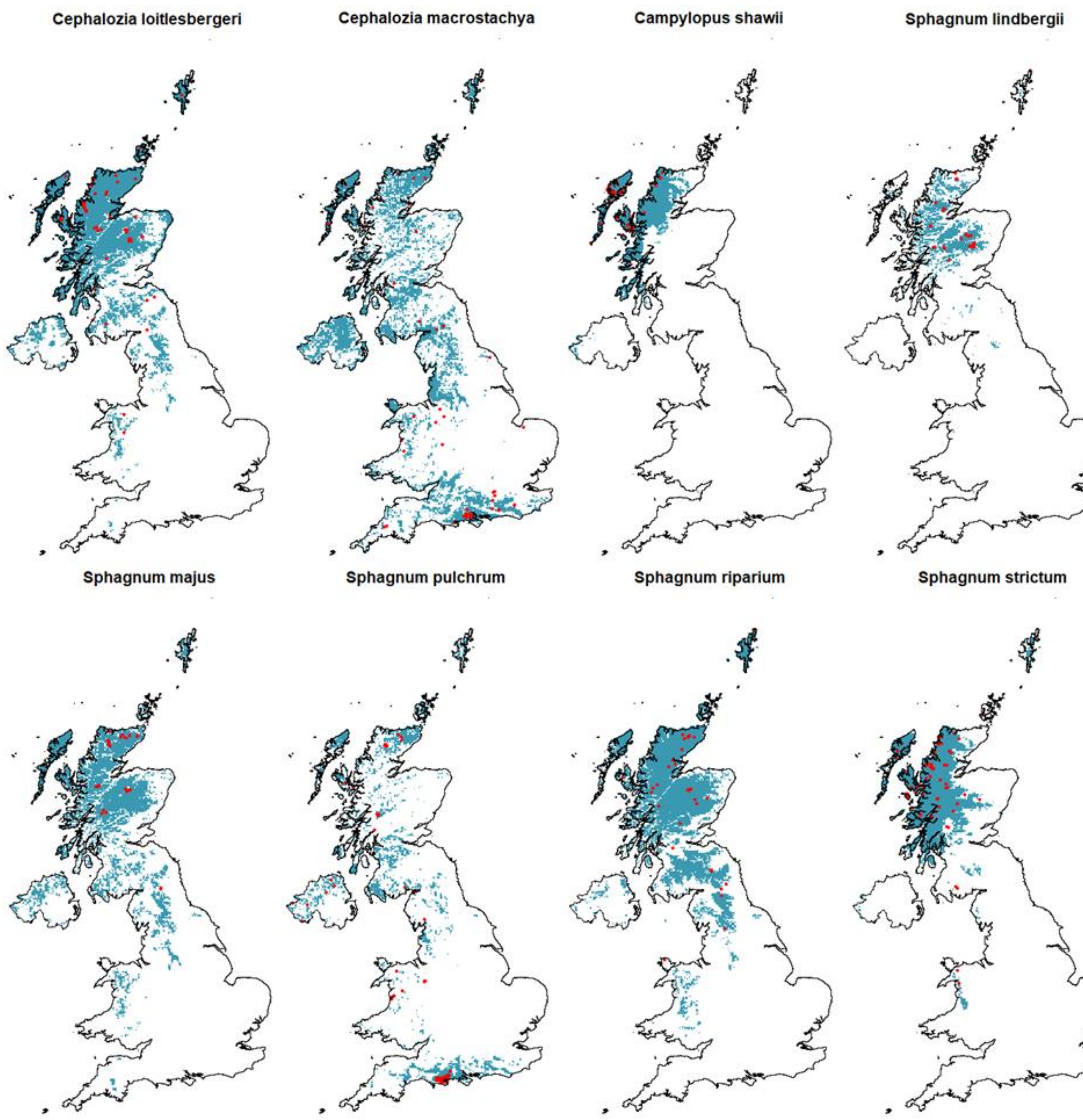


*Figure S6.2 Current species distribution modelled with Maxent in the UK (blue) and observed records (red points).*

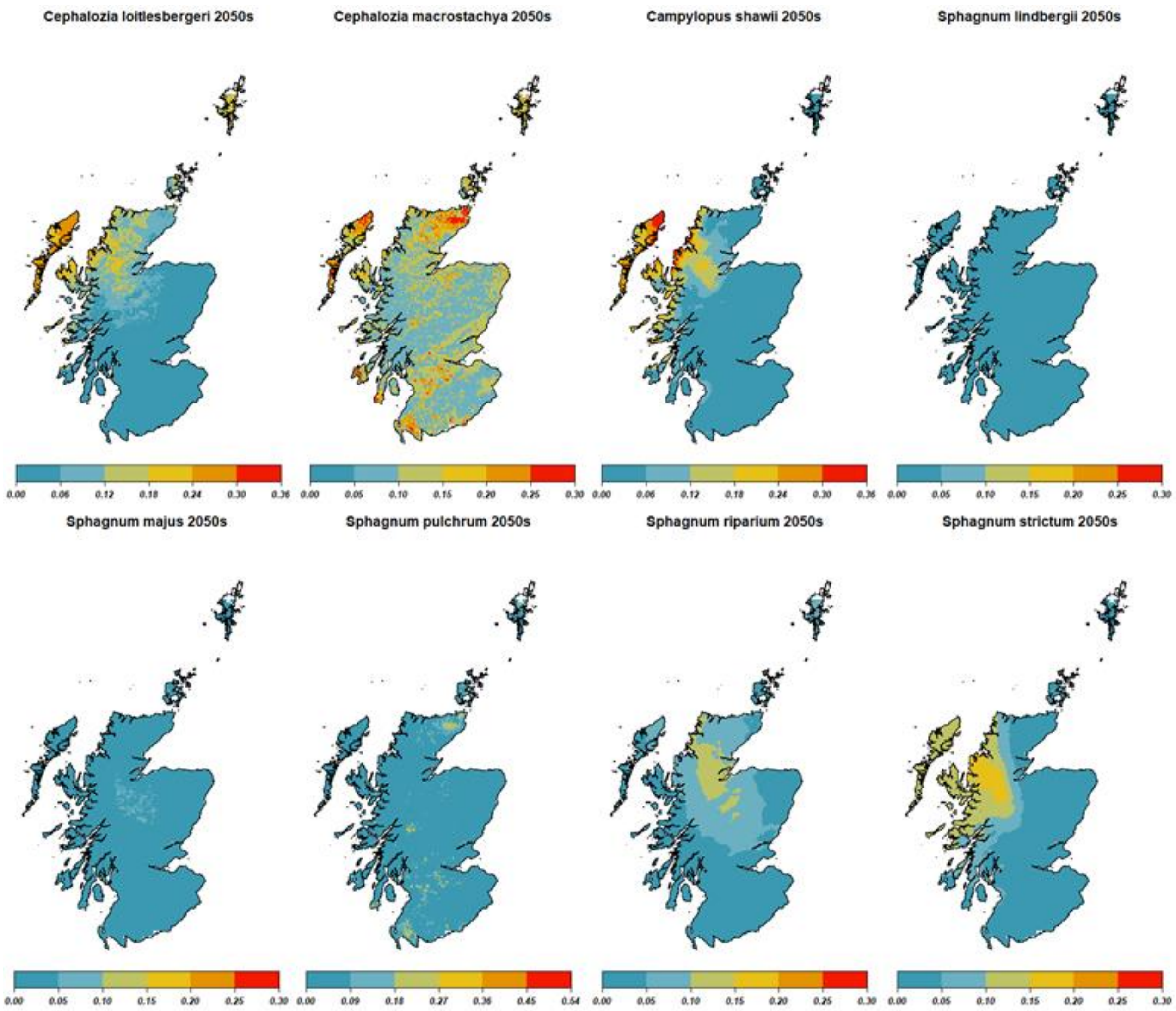


*Figure S6.3 Maxent future probability distribution for all the studied species of bryophytes ) obtained considering only changes in the climatic variables..*

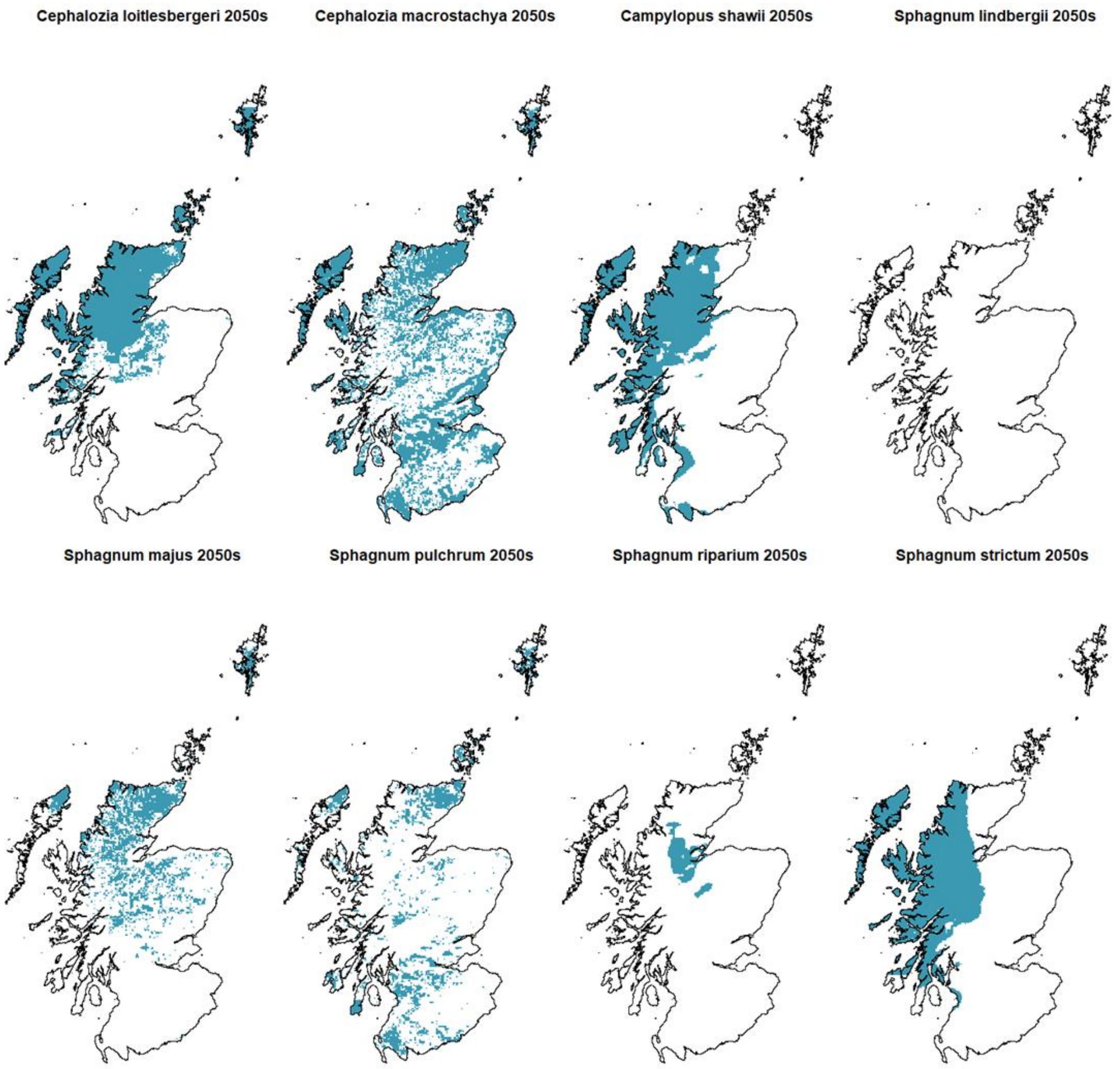


*Figure S6.4 2050s species distribution predicted with Maxent in Scotland (medium emission scenario) ) obtained considering only changes in the climatic variables.*

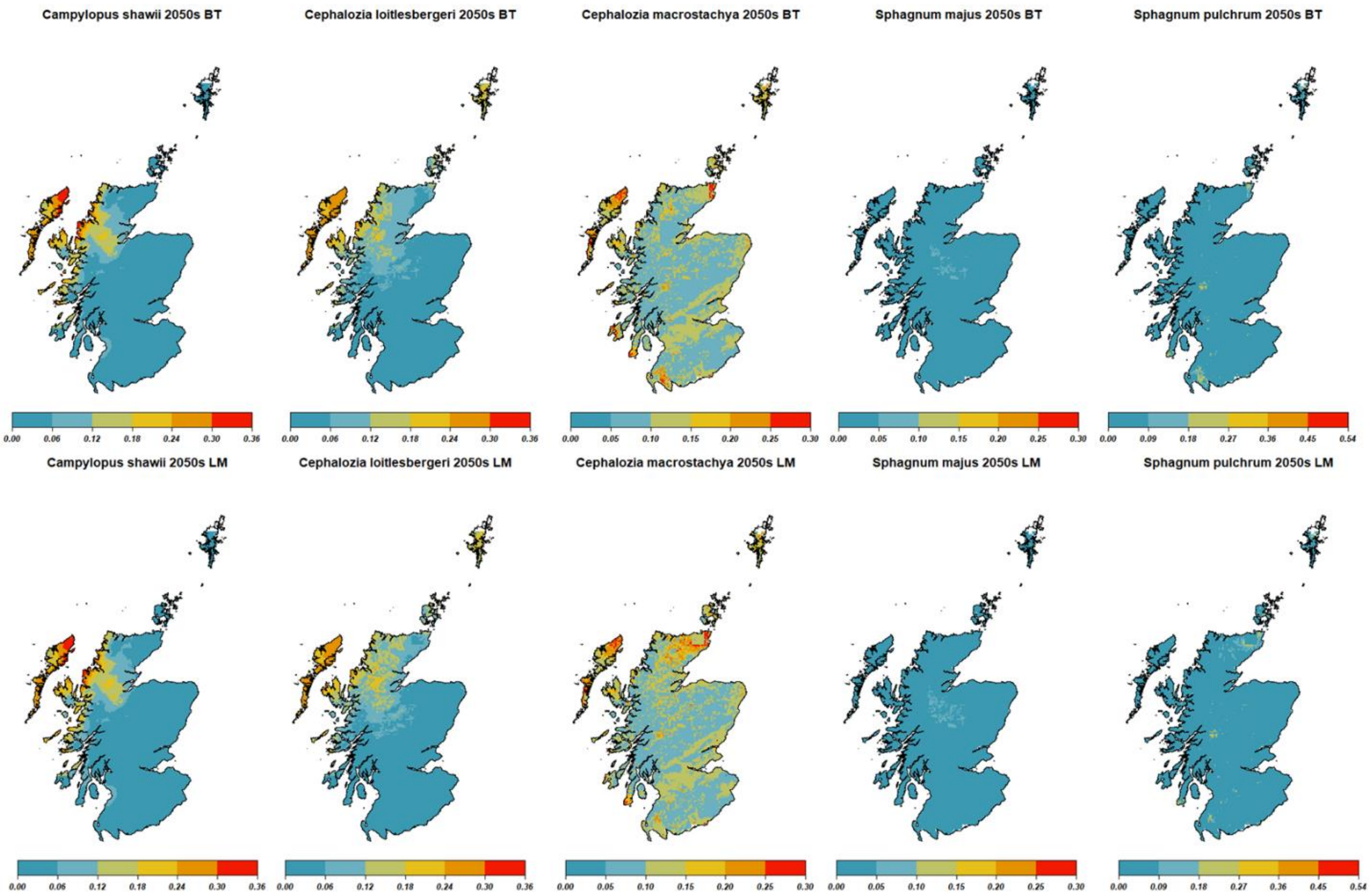


*Figure S6.5 Maxent future probability distribution for all the studied species of bryophytes using BT and LM blanket bog distribution*

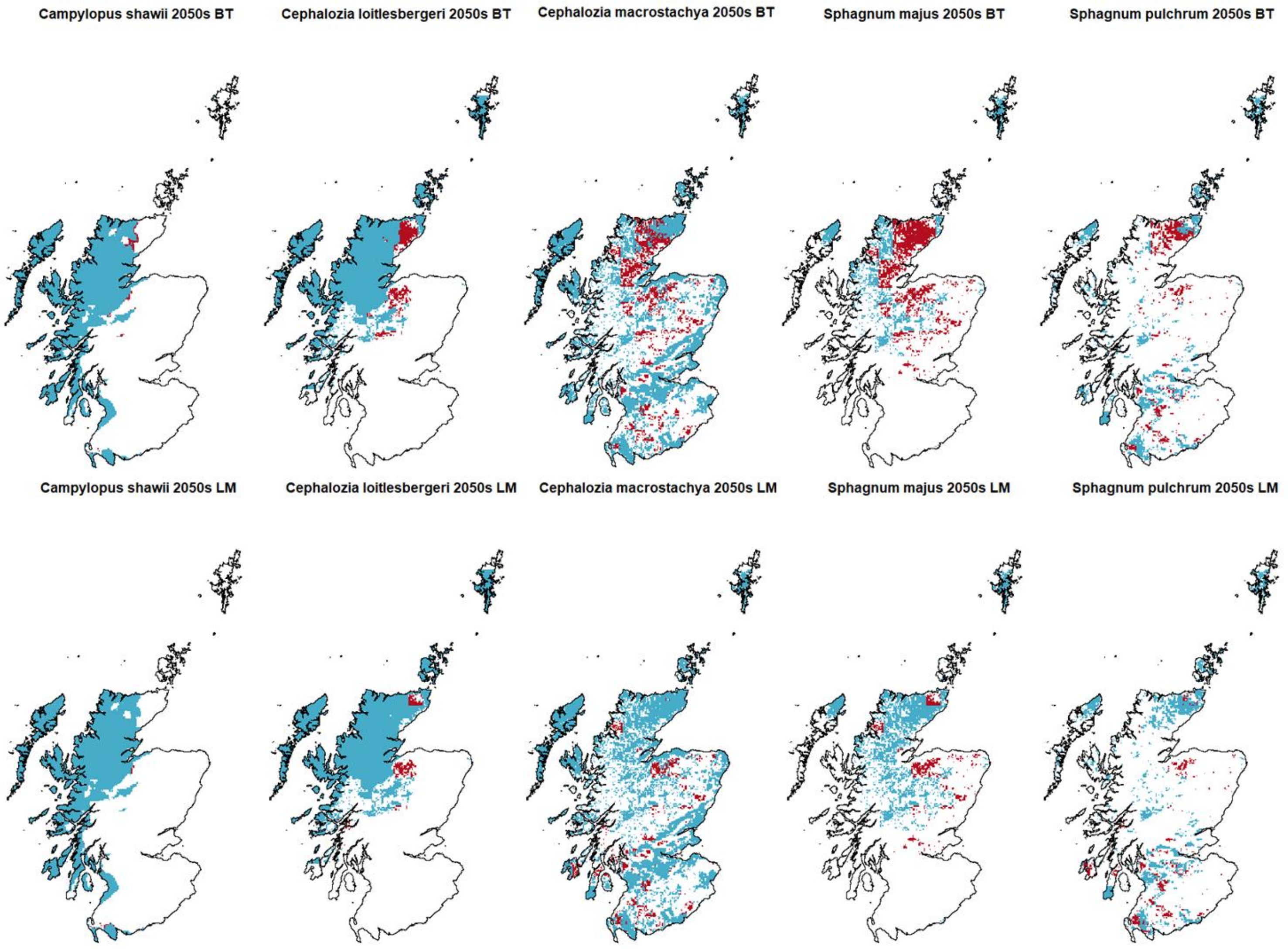


*Figure S6.6 Species distribution in the 2050s obtained with Maxent and using the BT (above) and LM (below) blanket bog distribution. In blue: future distribution as per the initial model (figure 3.2) that is unaffected by changes in the blanket bog layer. In red: areas that are predicted to be suitable by the initial model but are not suitable if blanket bog modelled change is considered.*

Figure S6.7 shows the changes in the future projections between the initial model run where the blanket bog layer has not been changed and the run with the BT and LM blanket bog distribution.

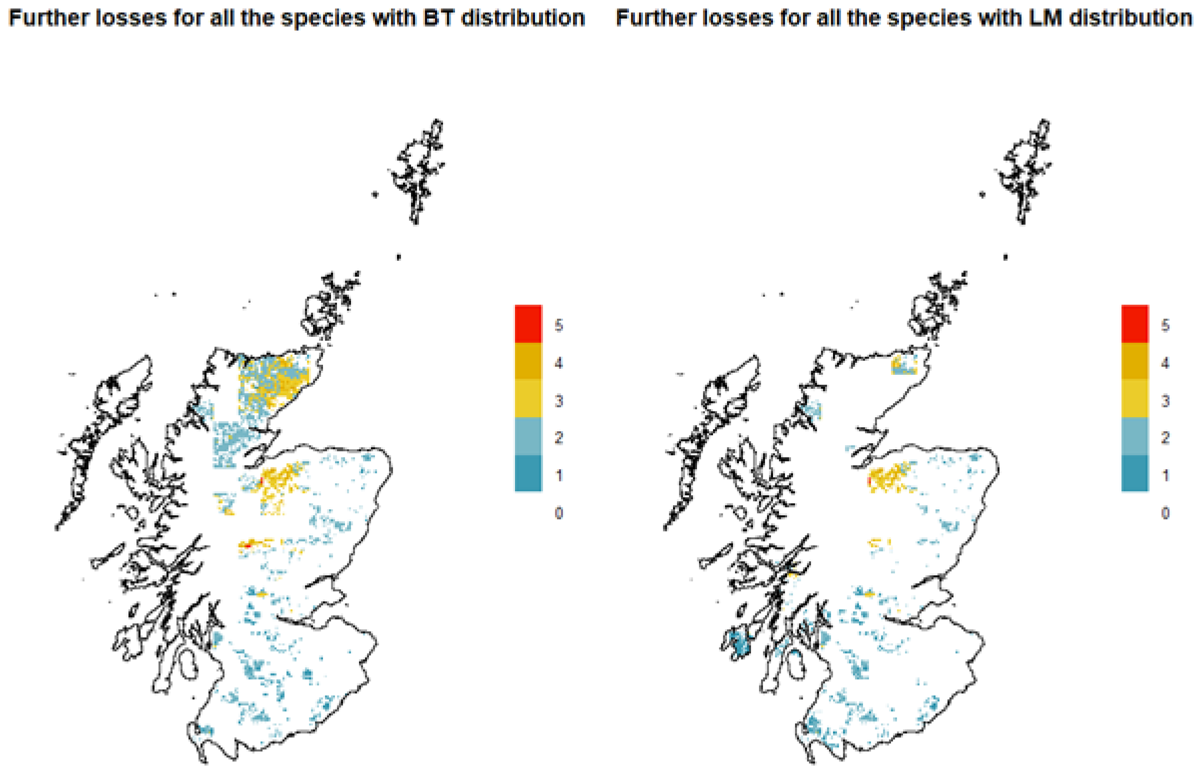


*Figure. S6.7 Areas most affected by the change in blanket bog modelled distribution, considering all the species (BT distribution on the left, LM distribution on the right). The legend shows the number of species involved.*

Figure S6.8 shows the results of the jackknife test, which indicates which variables contribute the most to increase the fit (the variable with the highest gain when used alone) and which variables carry more information by themselves (which lead to the highest loss of fit in the model if they are removed).

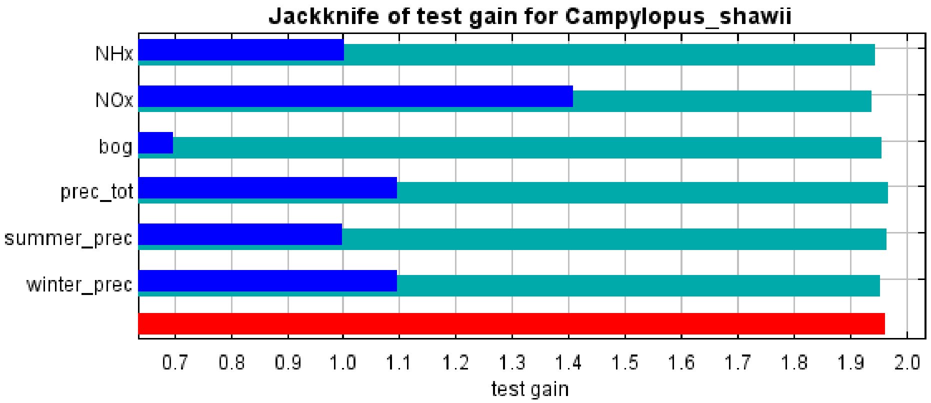


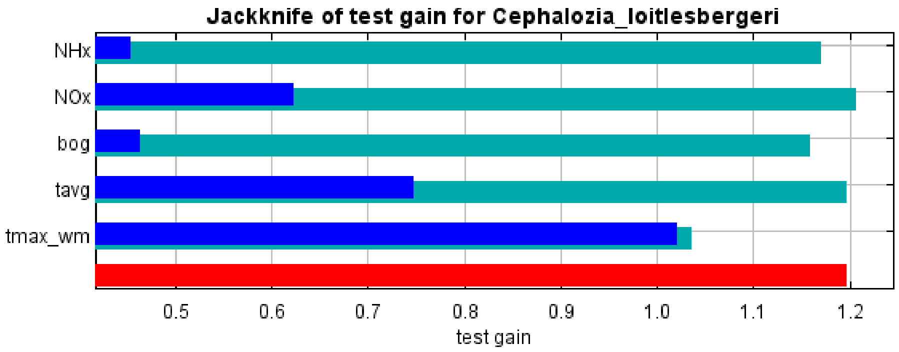


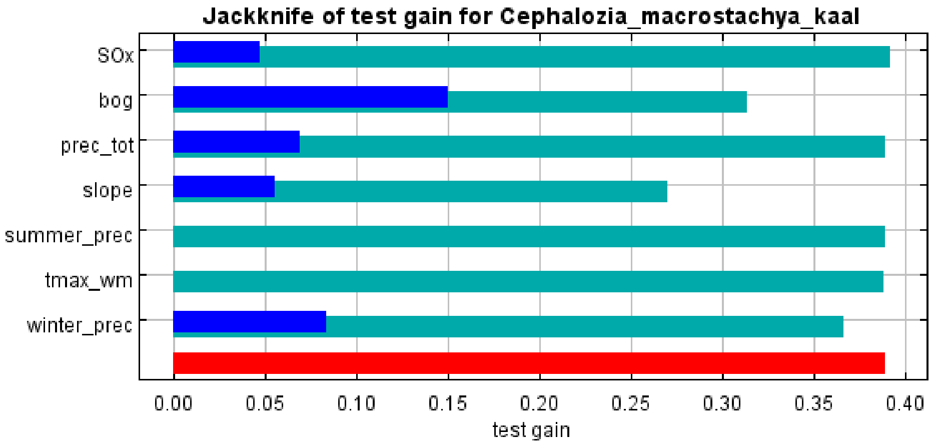


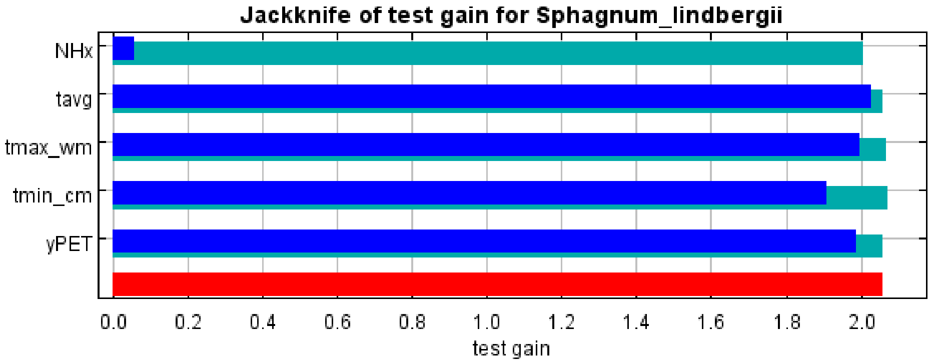

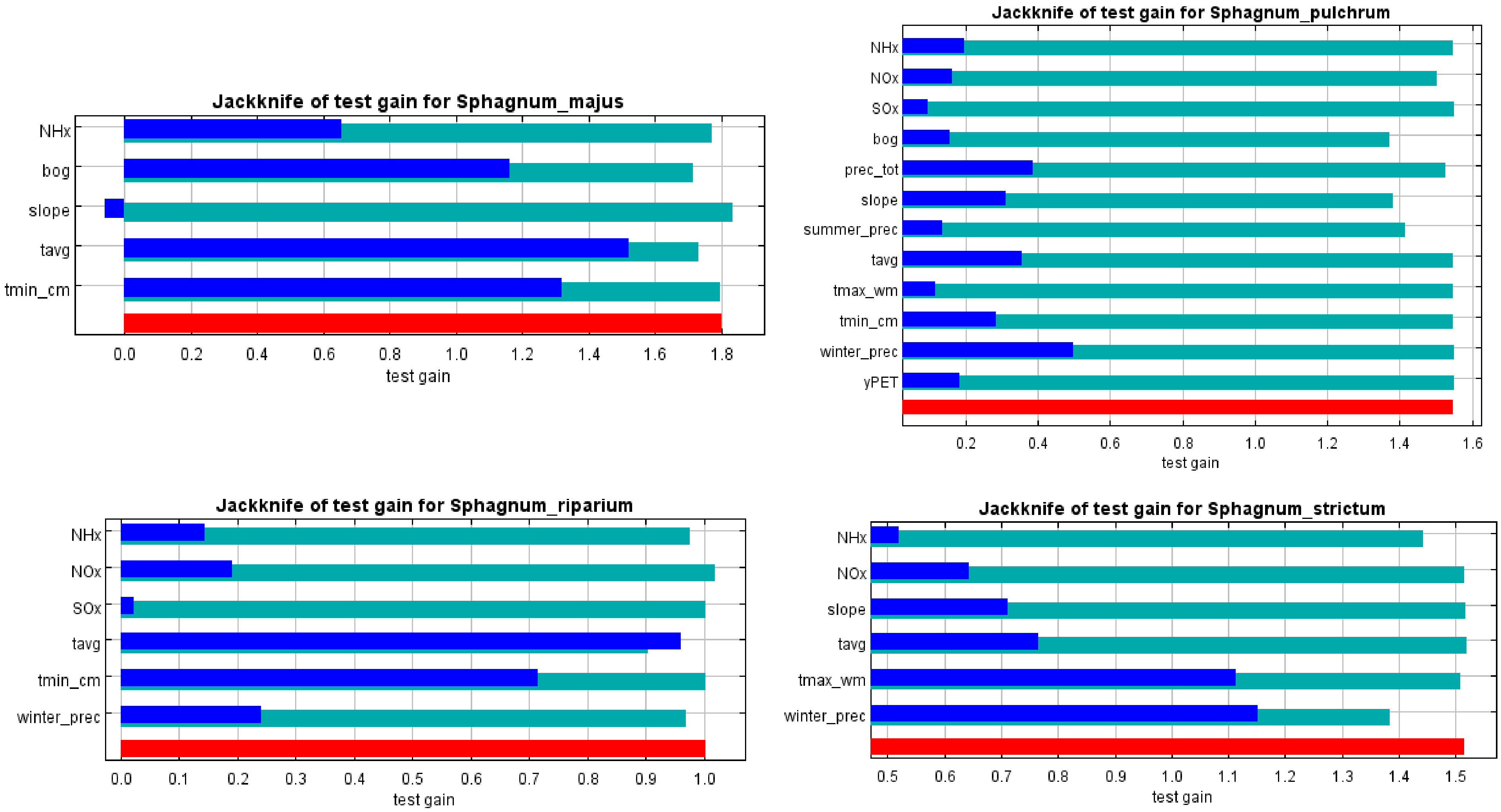


*Figure AVI.8 Outputs of the jackknife test produced by Maxent for each species. In light blue: test gain obtained by excluding the correspondent variable. In dark blue: test gain obtained by using only the correspondent variable. In red: test gain obtained including all the variables.*
